# Assessing the role of bacterial innate and adaptive immunity as barriers to conjugative plasmids

**DOI:** 10.1101/2024.04.12.588503

**Authors:** Berit Siedentop, Carlota Losa Mediavilla, Roger D. Kouyos, Sebastian Bonhoeffer, Hélène Chabas

## Abstract

Plasmids are ubiquitous mobile genetic elements, that can be either costly or beneficial for their bacterial host. In response to constant viral threat, bacteria have evolved various immune systems, such as the prevalent restriction modification (RM) (innate immunity) and CRISPR-Cas systems (adaptive immunity). At the molecular level, both systems also target plasmids, but the consequences of these interactions for plasmid spread are unclear. Using a modeling approach, we show that RM and CRISPR-Cas are effective as barriers against the spread of costly plasmids, but not against beneficial ones. Consequently, bacteria can profit from the adaptive advantages that beneficial plasmids confer even in the presence of bacterial immunity. While plasmids that are costly for bacteria may persist for a certain period in the bacterial population, RM and CRISPR-Cas pose a substantial burden for such plasmids, which can eventually drive them to extinction. Finally, we demonstrate that the selection pressure imposed by bacterial immunity on costly plasmids can be circumvented through a diversity of escape mechanisms and highlight how plasmid carriage might be common despite bacterial immunity. In summary, the population-level outcome of interactions between plasmids and defense systems in a bacterial population is closely tied to plasmid cost: Beneficial plasmids can persist at high prevalence in bacterial populations despite defense systems, while costly plasmids may face substantial reduction in prevalence or even extinction.

## 1 Introduction

In response to the selection pressure from bacteriophages, bacteria have evolved numerous immune systems, the two most frequent being Restriction-Modification (RM) systems and CRISPR-Cas systems [1]. Both systems provide protection by degrading phage dsDNA, but differ by their molecular mechanism. Briefly, RM-systems are typically composed of two main components: 1) a DNA-methylase and 2) a nuclease that cuts (i.e. restricts) unmethylated DNA at specific sequences, called restriction sites [2]. When non-methylated phage DNA enters the cell, there is a stochastic arms-race between the methylase and the nuclease. In the majority of cases, the nuclease detects this non-methylated (= non-self) DNA and restricts it before methylation occurs, excluding this foreign DNA. Rarely, however, methylation occurs before restriction, in which case the DNA gets recognized as self by the nuclease and remains in the cell. Because the immunity conferred by RM-systems is non-specific to a given phage, RM-systems are commonly referred as an innate bacterial immune system. On the contrary, due to the specificity of the immunity it confers, CRISPR-Cas systems are commonly referred as an adaptive immune system. Their specificity results from their molecular mechanism, which contains two main steps: spacer acquisition and interference. First, during the spacer acquisition step, some Cas proteins detect the entry of foreign DNA and pick up a 30-60bp DNA fragment called protospacer, which is then integrated into the CRISPR locus (where it is referred to as a spacer). Then, the CRISPR-Cas system protects against phage infection by using the spacers as RNA guides to bind the phage DNA by Watson-Crick pairing and trigger the degradation of the phage genome [3].

RM and CRISPR-Cas systems are both prevalent forms of bacterial immunity. Approximately 80% of bacteria carry at least one RM system and 40% of bacteria carry at least one CRISPR-Cas system [1] and on average a bacterial cell contains 2.3 RM systems and 0.52 CRISPR-Cas systems [1]. Within-cell heterogeneity, characterized by the presence of multiple RM systems per cell [1] or by the acquisition of several spacers targeting the same invader for CRISPR-Cas [4], contributes to a robust immune barrier [5]. Population-wide heterogeneity can also strengthen immunity. In the case of CRISPR-Cas, diversity in spacers across the population can efficiently eliminate a phage within a few days [6]. Concerning RM-systems, different strains within the population can carry different RM-systems, and while the role of population-based RM diversity is still to be better understood, it appears to confer advantages in phage defense [2, 7]. The widespread prevalence of these immune systems, along with the barriers they impose on phages, underscores their significance in anti-phage defense.

However, both CRISPR-Cas and RM systems are not exclusive to phages; they are also active at the molecular level against other mobile genetic elements, such as plasmids [8–11]. Plasmids are small dsDNA molecules and typically circular, that can spread both vertically and horizontally in bacterial populations. Horizontal transmission of conjugative plasmids happens through a mechanism called conjugation whereby the plasmid genome is transferred horizontally between bacterial cell as a ssDNA molecule. In contrast to virulent phages, plasmids often carry genes, like resistance genes, that can be beneficial to their bacterial hosts. Because of the molecular activity of both RM and CRISPR-Cas systems against plasmids, it was suggested that CRISPR-Cas and RM-systems could be a burden for the spread of plasmids and this could even be a disadvantage for bacteria if this prevents them from receiving plasmids with beneficial genes [12, 13].

There are several observations supporting the hypothesis that RM and CRISPR-Cas impose a burden on plasmids. First, in the lab, both RM-systems and CRISPR-Cas systems have been shown to decrease the conjugation/transformation rates of plasmids during short term assays [8, 14–16]. Second, bioinformatics analyses have shown that (i) in some bacterial species CRISPR-Cas presence is negatively correlated with antibiotic resistances [17, 18] and (ii) 38% of spacers found on chromosomal CRISPR locus target plasmids [19]. Third, recent findings indicate that plasmid genes exhibit a higher tendency to avoid restriction targets than chromosomal genes, suggesting that RM-systems impose a selection pressure on plasmids [20]. Fourth, both in the case of CRISPR-Cas and RM systems, plasmids can carry genes allowing circumvention of bacterial immunity, such as (i) anti-CRISPRs (small proteins inhibiting CRISPR-Cas interference) [21], (ii) anti-RMs (proteins inhibiting restriction) [22], (iii) Bet/Exo system (a system repairing the dsDNA break caused by CRISPR-Cas interference using a recombination mechanism) [23] or (iv) Toxin-Antitoxin (TA) systems (a system composed of a stable toxin and an unstable antitoxin, resulting in cell death after plasmid loss, for example because of CRISPR-Cas interference [24]). Additionally, it was shown that the plasmids’ leading region, the first region of the plasmid genome to enter the cell and be expressed [25], is enriched in anti-CRISPRs and anti-RMs. These genes are typically under the control of *Frpo* promoters, a family of promoters allowing transcription of ssDNA, enabling the expression of these genes before restriction and interference can occur [25], which suggest that their expression before the action of these immune systems is important for plasmid fitness. Finally, using experimental evolution and mathematical modeling, it was shown that bacteria carrying a type III CRISPR-Cas system targeting an antibiotic resistance plasmid lose their CRISPR-Cas immunity under strong antibiotic pressure in order to maintain the plasmid for survival [12].

Despite these lines of evidence, there are reasons to hypothesize that CRISPR-Cas and RM may not be a strong barrier for conjugative plasmids. First, the majority of studies reports a decrease in conjugation rate, not an absolute barrier [8, 14, 15], letting the question of the longer term dynamics unresolved. Second, even if in some bacterial species, there is an negative correlation between CRISPR-Cas and antibiotic resistances (as a proxy for plasmid carriage), global analyses don’t necessarily detect such a correlation and there are even reports of species for which there is a positive correlation between CRISPR-Cas and the carriage of antibiotic resistances [26, 27]. One could argue that the absence of negative correlation between CRISPR-Cas and antibiotic resistance genes results from the action of anti-CRISPRs. Yet, even though the role of anti-CRISPRs in these correlations have been rarely studied, a positive correlation between anti-CRISPRs and antibiotic-resistance genes was identified for several antibiotics in only one of the four pathogens studied [27]. Additionally, most of the experiments studying the evolutionary interactions between CRISPR-Cas and plasmids, have been carried with a bacterial population composed of cells already carrying a spacer targeting the plasmid [8, 12, 15], and sometimes using type III CRISPR-Cas system [12], which is known to be especially challenging to escape for phages [28]. While informative, these very specific conditions do not allow to explore the impact of the adaptive features of CRISPR-Cas, such as (i) spacer acquisition and (ii) population-based heterogeneity, on the evolutionary dynamics between CRISPR-Cas and plasmids. In the case of RM-systems, experimental studies do not consider that cells in a bacterial population carry different RM-systems, a feature that is predicted to be important for the understanding of phages-RM systems evolutionary dynamics [2, 7]. Therefore, our understanding of the dynamics between bacterial immunity and plasmids remain limited.

To understand if and how bacterial immunity affects the spread of plasmids, we model a bacterial population defending against a conjugative plasmid using either innate or adaptive immunity. Previous studies suggest that bacterial immunity can be lost in the presence of plasmids [12, 29]. Yet, bacteria exist in complex environments and face a high pressure due to virulent phages [30]. In this context, we assume that phages or other harmful mobile genetic element provide a selection pressure, preventing the loss of bacterial immunity. Hence, we explore potential plasmid spread in the presence of bacterial innate or adaptive immunity. We find that beneficial plasmids can usually spread, despite the presence of either innate or adaptive immunity. However, we find that both RM and CRISPR-Cas can impose a substantial burden on costly plasmids, potentially leading to their exclusion. This burden is not solely determined by the plasmid cost but can also be influenced by the strength of immunity and other plasmid fitness characteristics such as conjugation. Finally, we show that escape mechanisms such as Bet/Exo, TA systems and plasmid evolution can lower the burden imposed by bacterial immunity.

## 2 Methods: A mathematical framework to investigate the interaction between plasmids and bacterial innate and adaptive immunity

Here, we develop two mathematical models to explore the potential of RM and CRISPR-Cas to block plasmid spread (see figure 1.1 and SI sections 1.1 and 2.1). These models capture the following essential processes.

**Figure 1.**
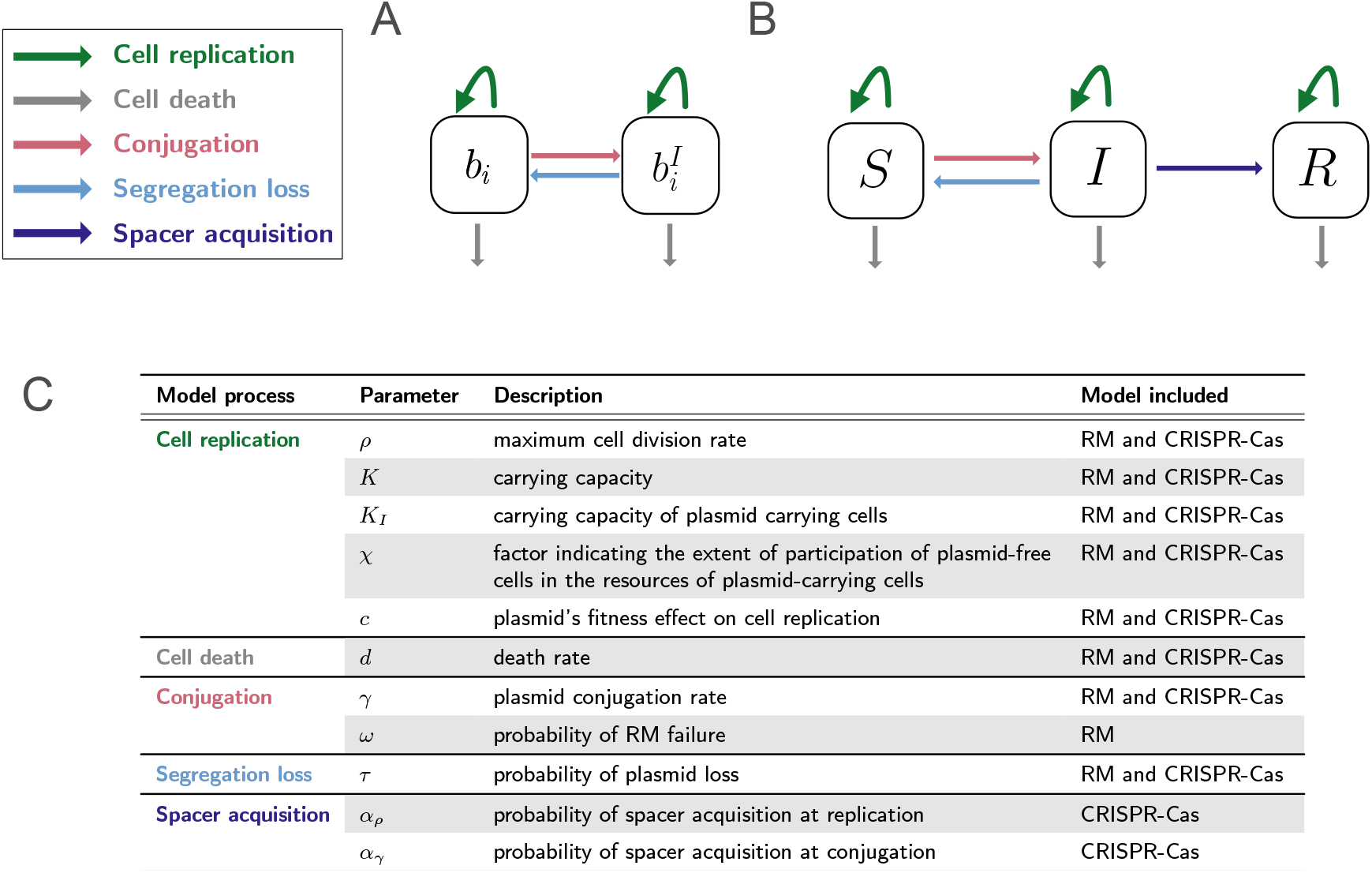
Visualization of modeled plasmid infection and immunity processes along with corresponding parameters. **A**. Schematic diagram of the basic RM model with detailed information available in supplement section 1.1. *b*_*i*_ represents bacterial cells with the RM systems set *i*, and 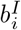 denotes cells infected with a plasmid, where the plasmid carries the corresponding methylation pattern. **B**. Schematic diagram of the basic CRISPR-Cas model with detailed information available in supplement section 2.1. *S* denotes cells with a naive to the plasmid CRISPR-Cas system, *I* bacterial cells with a naive CRISPR-Cas system that are infected with the plasmid, and *R* cells resistant to the plasmid. **C**. Plasmid infection and immunity processes are incorporated in the two basic immunity models, listed with their associated parameters, parameter descriptions, and indications of which model they occur in. The colors in the model process indicate the transition, as illustrated in (**A, B**), where the parameters are a driving factor. Choice of parameters is explained in SI sections 1.1.2, 2.1.1, and 1.1.4.

### Bacterial population composition

We consider a bacterial population defending either with CRISPR-Cas or with RM against a conjugative plasmid. We make the assumption that the bacterial population competes for and is limited by a single resource. In the case of RM-systems, the bacterial population consists of *n* strains, distinguished solely by the RM systems they carry. Each bacterial strain *b*_*i*_ carries a unique set *i* of RM-systems, where *i* ∈ 1, …, *n*. Importantly, we assume that the bacterial strains only differ in the set of RM-systems they carry and are identical otherwise. A bacterial cell *b*_*i*_ that gets infected with a conjugative plasmid with the corresponding methylation pattern is denoted by 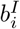 (figure 1A).

For CRISPR-Cas, we assume that the bacterial population consists of a clonal population with cells carrying a CRISPR-Cas system initially naive to the conjugative plasmid *S*. Once the naive bacterial cells are infected with the plasmid, they are denoted by *I*. If the CRISPR-Cas system acquires a spacer against the plasmid, making the cell immune, it is denoted as *R* (figure 1B).

### Bacterial population size

The bacterial growth is modeled by a density dependent division rate. For the bacterial population employing RM defense, the division rate is 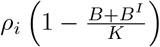, where *ρ*_*i*_ represents the maximal division rate of bacterial type *i, K* denotes the carrying capacity, and *B* and *B*^*I*^ are the total densities of plasmid-free and plasmid-carrying cells, respectively (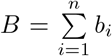 and 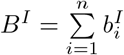). Importantly, in our simulations, we assume that the maximal division rate is the same for each bacterial type.

In some of our simulations, we assume that the conjugative plasmid can enable the use of a new energy source [31–33]. In our model, this is reflected by a higher carrying capacity *K*_*I*_ for bacterial cells carrying the plasmid than for those without (*K*_*I*_ *> K*). Plasmid-carrying cells can use the same resources as plasmid-free cells and an additional resource. The factor *χ* reflects how much plasmid-free cells *b*_*i*_ participate in the resources of plasmid-carrying cells 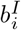. The density-dependent growth rate for plasmid-carrying cells is therefore described by 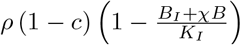, where *c* denotes the fitness effect of a plasmid on cell replication (*c >* 0: cost, *c <* 0: benefit).

Similarly, in the case of the population defending using CRISPR-Cas, the division rate for plasmid-free cells is given by 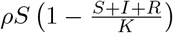, while for plasmid-carrying cells, it is expressed as 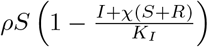. We assume a constant death rate *d* for all cells.

### Segregation loss and conjugation

Cell division of plasmid carrying cells may not necessary lead to two plasmid-carrying daughter cells. At cell division the plasmid gets lost from one daughter cell with probability *τ*, which is called segregation loss. We assume a well mixed bacterial population, where plasmid conjugation occurs between plasmid free cells and plasmid carrying in a density-dependent manner with conjugation rate *γ*.

### Bacterial immunity

In the context of the innate immunity the identity of donor and recipient is crucial for plasmid spread, i.e. which set of RM-systems *i* they carry. If the donor and recipient share the same set of RM-systems *i*, the conjugation rate is *γ* as the plasmid already possesses the corresponding methylation pattern. However, when the donor and recipient carry different sets of RM-Systems, the RM-System of the recipient acts as a barrier to successful plasmid conjugation. The incoming plasmid, with a methylation pattern recognized as foreign, is degraded. Nevertheless, RM-systems have a certain failure probability *ω*_*i*_, where the incoming plasmid may be methylated to the new host’s methylation pattern before being cleaved. Conjugation between bacterial cells of different types of RM-systems occurs therefore rarely with the rate *ω*_*i*_*γ*. In our simulations, we assume that each unique set of RM-systems has the same RM failure probability *ω* (see SI section 1.1.2 for further details).

In the context of CRISPR-Cas, immunity is modeled with spacer acquisition. Given that spacer acquisition has been linked to DNA replication [10], we assume that spacer acquisition from the plasmid can occur whenever the plasmid is replicated, i.e., during both cell division and conjugation. We denote the probability of acquiring a spacer at cell division as *α*_*ρ*_ and the probability of acquiring a spacer at conjugation as *α*_*γ*_ (see SI section 2.1.1 for further details).

### Implementation

For our investigations, we initiated simulations in both the RM and CRISPR-Cas models with a bacterial population that has reached equilibrium population size, with only a small fraction of plasmid-carrying cells. In the case of RM-systems, we assumed that only a small fraction of one type of bacteria *b*_*i*_ is initially infected with the plasmid, and the rest of the population consists of plasmid-free cell types distributed in equal proportion throughout the bacterial population. In the case of CRISPR-Cas, we assumed that no cell has acquired a spacer against the plasmid yet. Unless specified otherwise, we assume that the plasmid does not confer the ability to utilize an additional resource, implying *K* = *K*_*I*_ and *χ* = 1 (see SI sections 1.1.3 and 2.1.2 for further details).

The simulations for RM were run stochastically due to potential qualitative differences from deterministic implementations, particularly for costly plasmids (see SI section 1.1.3). The probability of plasmid extinction was calculated for each simulated parameter set based on 100 simulations. For stochastic simulations, we used the *adaptivetau* package (version 2.2-3) in R (version 4.2.2), implementing the Gillespie algorithm with adaptive *tau*-leaping [34, 35]. The simulations for the basic CRISPR-Cas model were implemented deterministically with JULIA (version 1.8.5) and the *DifferentialEquations* package (verison 7.7.0) [36]. Global sensitivity analysis was performed with the Sobol’s method, a variance based sensitivity analysis, implemented in the JULIA package *GlobalSensitivity* (version 2.1.4) [37] (more details in SI sections 2.1.7, 2.4.2). The more complex CRISPR-Cas model, taking CRISPR-Cas and plasmid co-evolution into account (SI section 2.3), is implemented in R stochastically using the *adaptivetau* package.

## 3 Results

We investigate the spread of plasmids in a bacterial population at maximal population size, starting with a small fraction of plasmid-carrying cells, in the presence of CRISPR-Cas and RM-systems. Our results are structured in three parts. First, using our two mathematical models of RM and CRISPR-Cas, we explore the potential spread of a beneficial plasmid within a bacterial population in the presence of bacterial immunity. Second, we assess the extent to which RM and CRISPR-Cas serve as barriers for costly plasmids. Third, we evaluate mechanisms that can allow costly plasmids to reduce the burden of bacterial immunity.

### 3.1 Bacterial immune systems let beneficial plasmids spread in bacterial populations

Our models predict that plasmids with a positive fitness effect can successfully invade and persist in bacterial populations despite immunity from RM-systems or CRISPR-Cas (figure 2). Overall, our simulations demonstrate mostly robustness in this outcome to variations in plasmid characteristics and the strength of bacterial immunity (SI sections 1.1.4, 2.1.4). In rare cases, CRISPR-Cas might exclude beneficial plasmids if the benefit is very low (SI sections 2.1.3, 2.1.4). In both cases of bacterial immunity, RM-systems or CRISPR-Cas, bacterial growth competition ensures plasmid maintenance, as explained in detail below.

**Figure 2.**
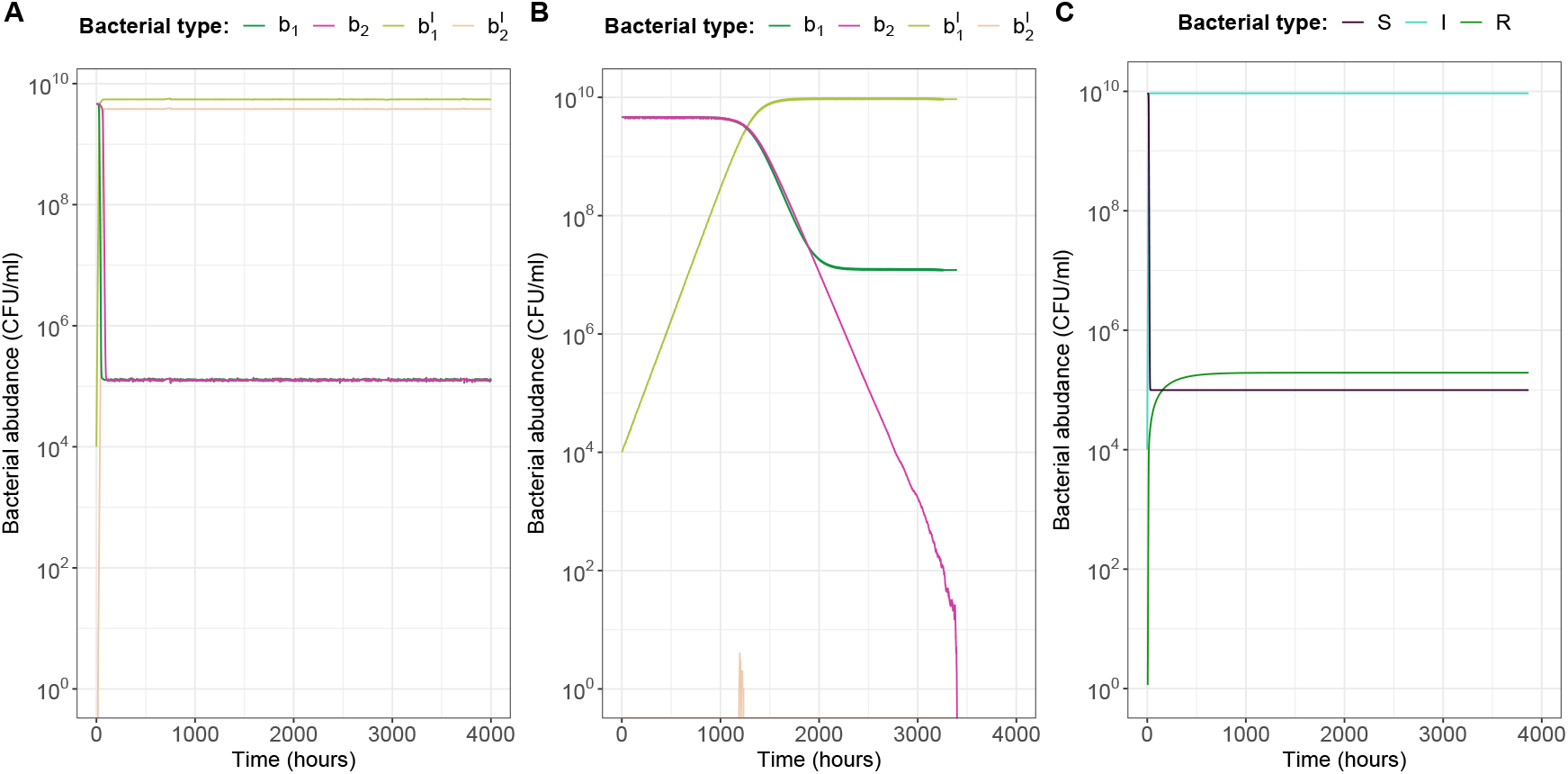
Beneficial plasmids can spread despite RM-systems (A,B) and CRISPR-Cas (C). **A** and **B** are two exemplary simulations for RM-systems showing the two distinct scenarios of maintenance of beneficial plasmids: **A** where the beneficial plasmid is able to spread to each individual strain in the population as it overcomes all RM-system barriers, and **B** where the beneficial plasmid can only spread in the population for which the plasmid has the corresponding methylation pattern, unable to overcome other strain’s RM-system. The population carrying the plasmid out-competes plasmid-free populations due to the growth benefit conferred by the plasmid. Plasmid-free cells are denoted by *b*_*i*_ and plasmid carrying cells by 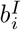, where *i* indicates the identify of the strain. Both simulations are run with a beneficial plasmid fitness effect *c* = − 0.1 and a RM-systems failure probability of *ω* = 10^−8^. The plasmid conjugation rate is set to *γ* = 10^−10^ and *γ* = 10^−13^ respectively. **C** Exemplary simulation of the spread of beneficial plasmids in a population carrying an initially naive CRISPR-Cas system. Cells carrying a naive CRISPR-Cas system are denoted by *S*, cells carrying a naive CRISPR-Cas system and infected by a plasmid are denoted by *I*, and cells that acquired a spacer and are resistant against the plasmid are denoted by *R*. The conjugation rate is set to *γ* = 10^−10^ and the probability of acquiring a spacer at conjugation and replication is set to *α*_*ρ*_, *α*_*γ*_ = 10^−6^. The probability of losing a plasmid was set to *τ* = 10^−4^ in all three simulations.

In the RM-model, the beneficial plasmid initially spreads within the bacterial sub-population carrying the corresponding methylation pattern. While the beneficial plasmid persists in this population, there are two potential outcomes: either the plasmid spreads to other bacterial strains (figure 2A), or the plasmid-carrying strain out-competes others due to the plasmid’s failure to overcome the barriers imposed by RM-systems in time (figure 2B). The likelihood of each outcome depends on the plasmid’s ability to overcome barriers of distinct RM-systems in plasmid-free bacteria before being out-competed by already plasmid-carrying bacteria. Factors like the plasmid conjugation rate, the growth advantage conferred by the plasmid, and the probability of failure of the RM-system determine the occurrence of these scenarios (SI section 1.1.5, figure S3). However, the overall pattern is robust that plasmids with positive fitness effect are maintained (figure S2).

In the CRISPR-Cas model, a beneficial plasmid initially spreads within the naive population. Although CRISPR-Cas occasionally acquires spacers, resulting in the production of plasmid resistant cells, these resistant cells persist at low frequencies (figure 2C). This is because they are out-competed by naive cells carrying the plasmid, which have a higher growth rate. This point can be illustrated by the analytical solution of a specific scenario of the CRISPR-Cas model. Assuming all naive CRISPR cells are infected by plasmids and the plasmid has no segregation loss, stable plasmid maintenance occurs if the plasmid fitness advantage compensates for spacer acquisition (SI section 2.1.3). Given that the known estimates of spacer acquisition, derived from phages, are rather low, around 10^−6^ [38], plasmids with relatively low growth benefits can still maintain in the population despite the presence of CRISPR-Cas immunity. Rarely, when the plasmid benefit is too low, plasmid may not persist (SI sections 2.1.3, 2.1.4). Simulations using the full CRISPR-Cas model show that the benefits of beneficial plasmids not only allow them to invade and survive in initially naive populations but also in populations with a high proportion of resistant cells (SI figure S7).

In the main text, we only present results for beneficial plasmids that increase the maximal bacterial growth rate. We observe similar results for plasmids that enable the utilization of an additional resource and thereby increase the maximal population size (SI sections 1.1.6 and 2.1.6). Overall, we conclude that beneficial plasmids are able to maintain in bacterial populations despite bacterial immunity. However, especially in the case of RM-systems, bacterial strains can face a disadvantage if their barrier to plasmid entry is too high, resulting in reduced RM-diversity within the population (figure 2B, SI section 1.1.5).

### 3.2 The strength of bacterial immunity and plasmid fitness determine the blocking of costly plasmids

In contrast to their impact on beneficial plasmids, we find that both RM-systems and CRISPR-Cas can block costly plasmids, but the infection dynamics between the two types of bacterial immune systems differ.

RM-Systems act as an immediate barrier. However, once a plasmid acquires the appropriate methylation pattern, it is recognized as ‘self,’ rendering RM-Systems ineffective in blocking further plasmid spread. In our model, a costly plasmid can only be maintained long-term if it overcomes each bacterial strain’s RM barrier in time before plasmid-free strains out-compete plasmid-carrying cells due to the plasmid’s growth disadvantage. Failure to overcome a single strain’s RM barrier leads to the plasmid’s extinction. Persistence of a costly plasmid is determined by both RM and plasmid parameters. Two parameters in our RM-model determine the strength of RM-immunity: first, the probability of RM failure, and second, the diversity of distinct RM-systems in the bacterial population. A low failure rate and a high diversity of RM-systems within the bacterial population make it difficult for the plasmid to spread to all bacterial strains before being out-competed by plasmid-free bacteria (figure 3C). On the plasmid side, a higher conjugation rate and lower plasmid cost increase the likelihood of plasmid persistence despite RM-immunity (figure 3A,B). A higher conjugation rate provides more opportunities for the plasmid to overcome RM-system barriers, while lower plasmid cost allows more time for overcoming all RM-system barriers before being out-competed.

**Figure 3.**
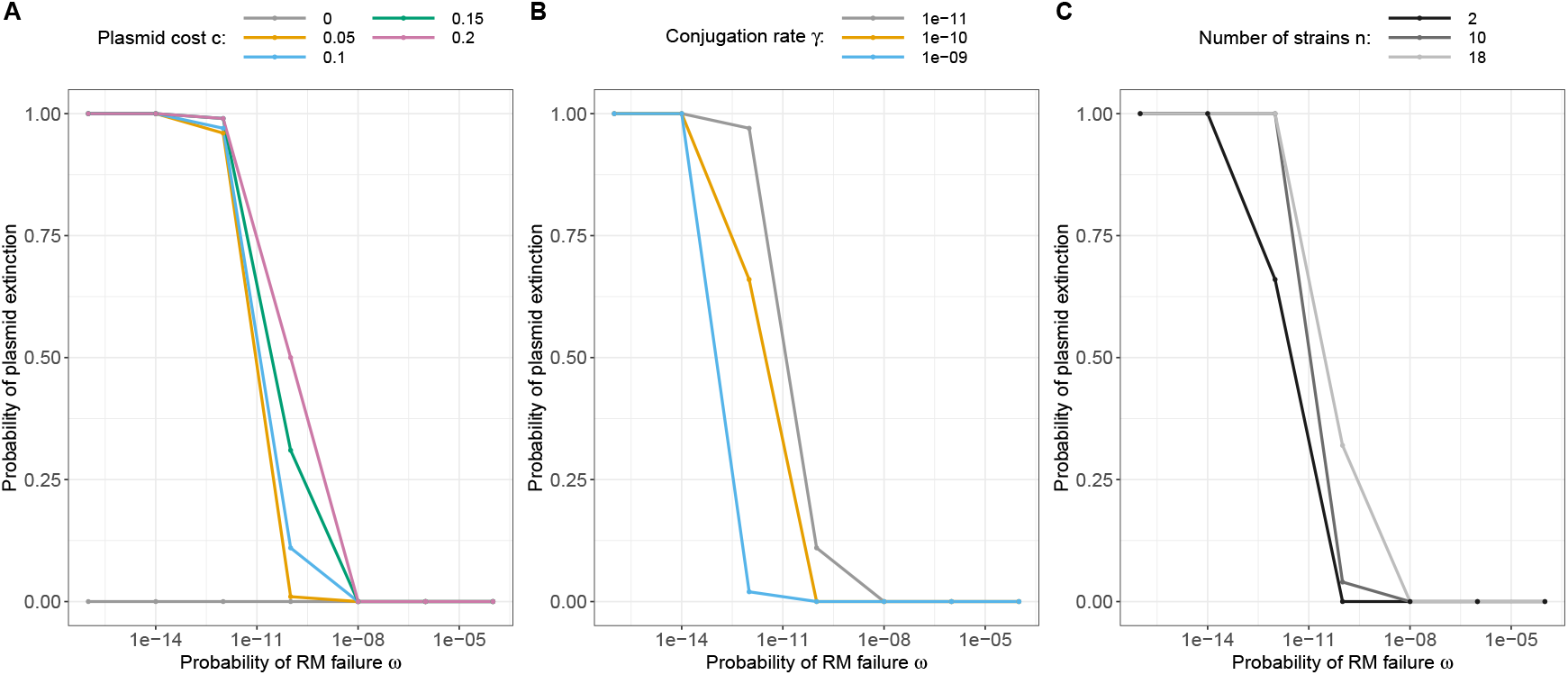
Impact of plasmid fitness and strength of RM-systems on plasmid extinction probability. The figure shows the probability of plasmid extinction based on 100 stochastic simulations. The dependencies include the probability of RM failure *ω* and (**A**) plasmid fitness *c*, (**B**) conjugation rate *γ*, or (**C**) the number of distinct bacterial strains n. The probability of segregation loss is set to *τ* = 10^−4^ in all simulations, plasmid fitness effect is set to *c* = 0.1 in **B,C**, and the conjugation rate is set to *γ* = 10^−11^ in **A,C**.

CRISPR-Cas defence does not rely on pre-existing immunity; instead, immunity can be acquired during the infection process. Our simulations show that costly plasmids are able to spread initially in the naive population. However, due to the plasmid’s fitness cost, the acquired resistant population gains a growth advantage, leading to long-term out-competition of plasmid-carrying cells. Our simulations show that the time until the plasmid is eliminated can be long (figure 4A). Global sensitivity analysis indicates that the speed with which the plasmid is eliminated is highly dependent on the cost the plasmid confers and only weakly depends on the probability of spacer acquisition (figure 4B). This can be explained by the fact that spacer acquisition is crucial for founding the resistant population, but the increase in the resistant population is predominantly driven by higher growth fitness in comparison to the plasmid-carrying cells rather than through new spacer acquisition. However, when extending our basic CRISPR-Cas model (SI section 2.3) to account for certain complexities in the co-evolution between CRISPR-Cas and plasmids — such as plasmid spacer escape and population-wide spacer heterogeneity — we find that the strength of CRISPR-Cas immunity, represented in our model by the probability of spacer acquisition, plays a more crucial role. In this complex model, the presence of spacer acquisition not only matters, as in the basic CRISPR-Cas model, but the magnitude of the probability of spacer acquisition substantially influences the reduction in plasmid prevalence (figure 4C). Further discussion on co-evolution between CRISPR-Cas and plasmids is provided later on.

**Figure 4.**
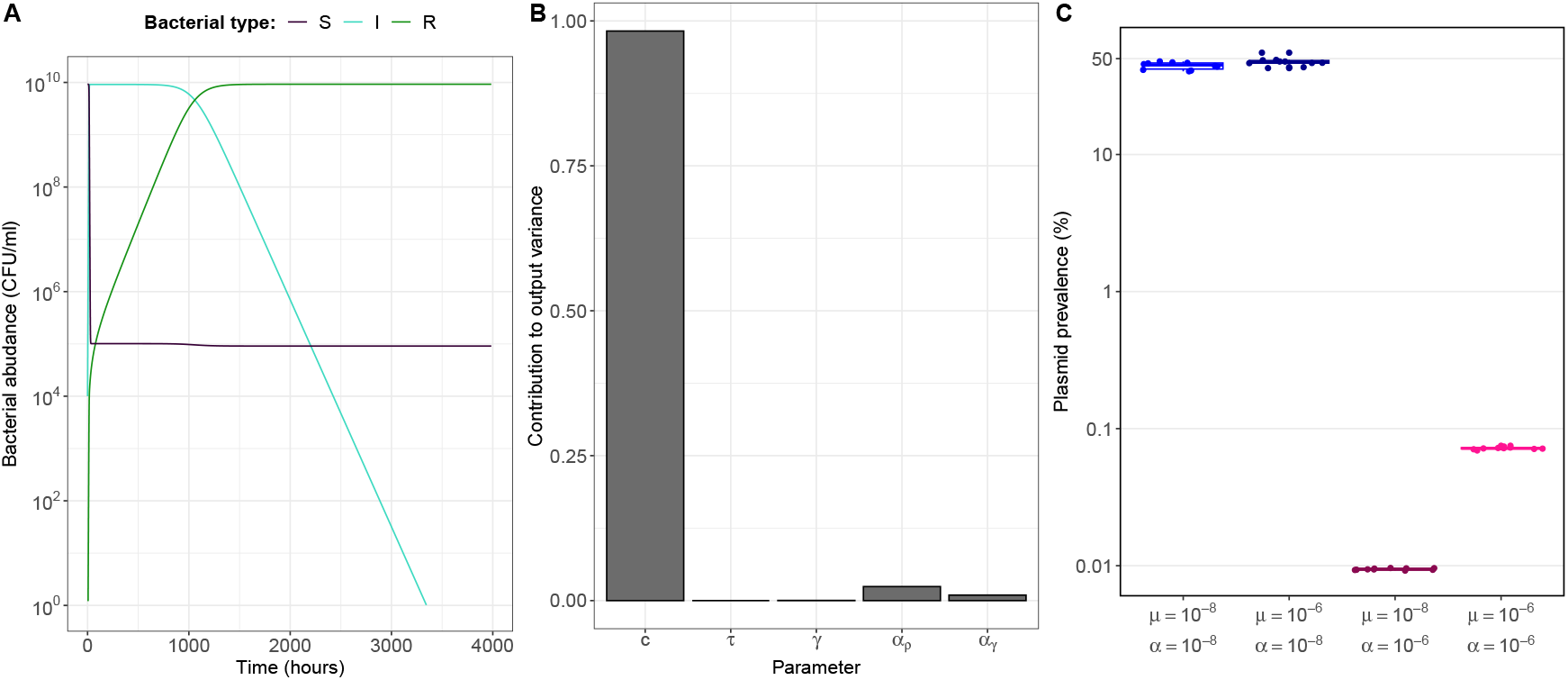
Impact of plasmid fitness and the strength of CRISPR-Cas on plasmid elimination. (**A**) Exemplary simulation showing the infection dynamics of a costly plasmid with fitness effect *c* = 0.1 in an bacterial population with an initially naive CRISPR-Cas system. Conjugation rate is set to *γ* = 10^−10^, probability of segregation loss *τ* = 10^−4^, probability of acquiring a spacer at replication and conjugation *α*_*ρ*_, *α*_*γ*_ = 10^−6^. (**B**) Total order Sobel indices indicating the importance of parameters to explain how long a costly plasmid takes to be eliminated from the bacterial population. *c* is the plasmid fitness effect, *τ* the probability of segregation loss, *γ* the conjugation rate, *α*_*ρ*_ the probability to acquire a spacer at cell division and *α*_*γ*_ the probability to acquire a spacer at conjugation. Note that total order Sobol indices also includes interactions of parameters and therefore does not necessarily add up to one. See SI section 2.1.7 for a detailed explanation and parameter bounds used for the global sensitivity analysis. **C** Plasmid prevalence with varying mutation probability of a plasmid protospacer *μ* and *α* probability of acquiring a spacer. Both these events are assumed to happen at replication and conjugation with the same probability. We show 10 simulations per parameter set and the corresponding boxplot, further detail is given in SI section 2.3)

In conclusion, the spread of costly plasmids in bacterial populations defended by RM-systems or CRISPR-Cas depends on the strength of these immune systems and plasmid fitness. Crucial factors for RM-system immunity strength include the probability of RM-system failure and RM-diversity within the bacterial population. For CRISPR-Cas, strength is determined by the probability of spacer acquisition. Additionally, plasmid cost substantially influences the dynamics of plasmid spread for both RM-systems and CRISPR-Cas. Moreover, we demonstrate that higher conjugation increases the likelihood of maintaining costly plasmids in bacterial populations defended by RM-systems.

### 3.3 Mechanisms allowing evasion of bacterial immunity

While RM-systems and CRISPR-Cas impose barriers for costly plasmids, several mechanisms exist to alleviate the burden of bacterial immunity for conjugative plasmids (table 1). In the discussion section, we provide an overview and evaluation of escape mechanisms, including commonly found anti-RM and anti-CRISPR proteins. In the following we investigate three specific mechanisms in more detail: TA-systems, the *Bet/Exo* system, and evasion of CRISPR-Cas spacers through plasmid evolution.

**Table 1.**
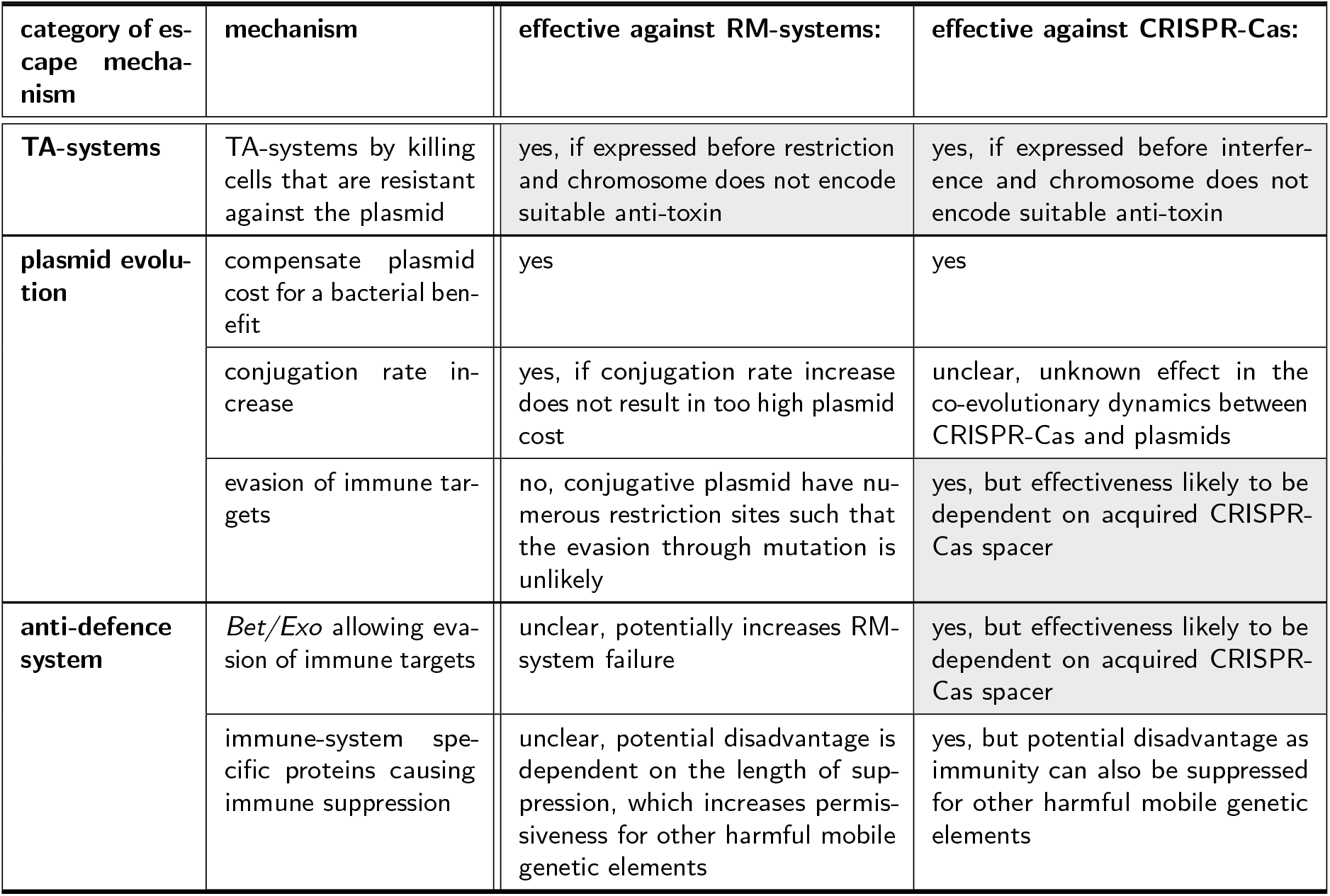
Summary and assessment of mechanisms lowering the burden of RM-systems and CRISPR-Cas for plasmids. Fields highlighted in grey indicate mechanisms presented in detail in the result section. A discussion of these and additional mechanisms can be found in the discussion section.

| category of escape mechanism | mechanism | effective against RM-systems: | effective against CRISPR-Cas: |
| --- | --- | --- | --- |
| **TA-systems** | TA-systems by killing cells that are resistant against the plasmid | yes, if expressed before restriction and chromosome does not encode suitable anti-toxin | yes, if expressed before interference and chromosome does not encode suitable anti-toxin |
| **plasmid evolution** | compensate plasmid cost for a bacterial benefit | yes | yes |
|  | conjugation rate increase | yes, if conjugation rate increase does not result in too high plasmid cost | unclear, unknown effect in the co-evolutionary dynamics between CRISPR-Cas and plasmids |
|  | evasion of immune targets | no, conjugative plasmid have numerous restriction sites such that the evasion through mutation is unlikely | yes, but effectiveness likely to be dependent on acquired CRISPR-Cas spacer |
| **anti-defence system** | *Bet/Exo* allowing evasion of immune targets | unclear, potentially increases RM-system failure | yes, but effectiveness likely to be dependent on acquired CRISPR-Cas spacer |
|  | immune-system specific proteins causing immune suppression | unclear, potential disadvantage is dependent on the length of suppression, which increases permissiveness for other harmful mobile genetic elements | yes, but potential disadvantage as immunity can also be suppressed for other harmful mobile genetic elements |

#### Toxin-Antitoxin systems

Plasmid commonly carry Toxin-Antitoxin (TA) systems. When incorporated into our basic CRISPR-Cas model (SI section 2.2), our model illustrates that TA-system, if active before CRISPR-Cas interference, can support plasmid survival in two non-mutually exclusive ways: (i) preventing the emergence of resistant cells and (ii) killing resistant cells through plasmid conjugation. In our basic CRISPR-Cas model, when a cell acquires a spacer and the TA-system is expressed beforehand, resulting in the death of newly resistant cells, it hinders the establishment of the resistant population. However, if we account for the possibility of TA-system failure or occasional interference by CRISPR-Cas before TA-system expression, a small fraction of resistant cells survives, allowing the resistant population to establish (figure 5A). When considering that conjugation to resistant cells results in cell death for a fraction of these recipients due to toxins produced before plasmid degradation, we observe the persistence of costly plasmids over time. The survival of the plasmid through killing resistant cells by conjugation is also more robust to the failure of the TA-system or occasional CRISPR-Cas interference before TA-system expression (figure 5B). Our results thus support the hypothesis proposed by Van Sluijs et al. that expressing plasmid-encoded toxins before CRISPR-Cas interference can explain plasmid survival despite CRISPR-Cas [39].

**Figure 5.**
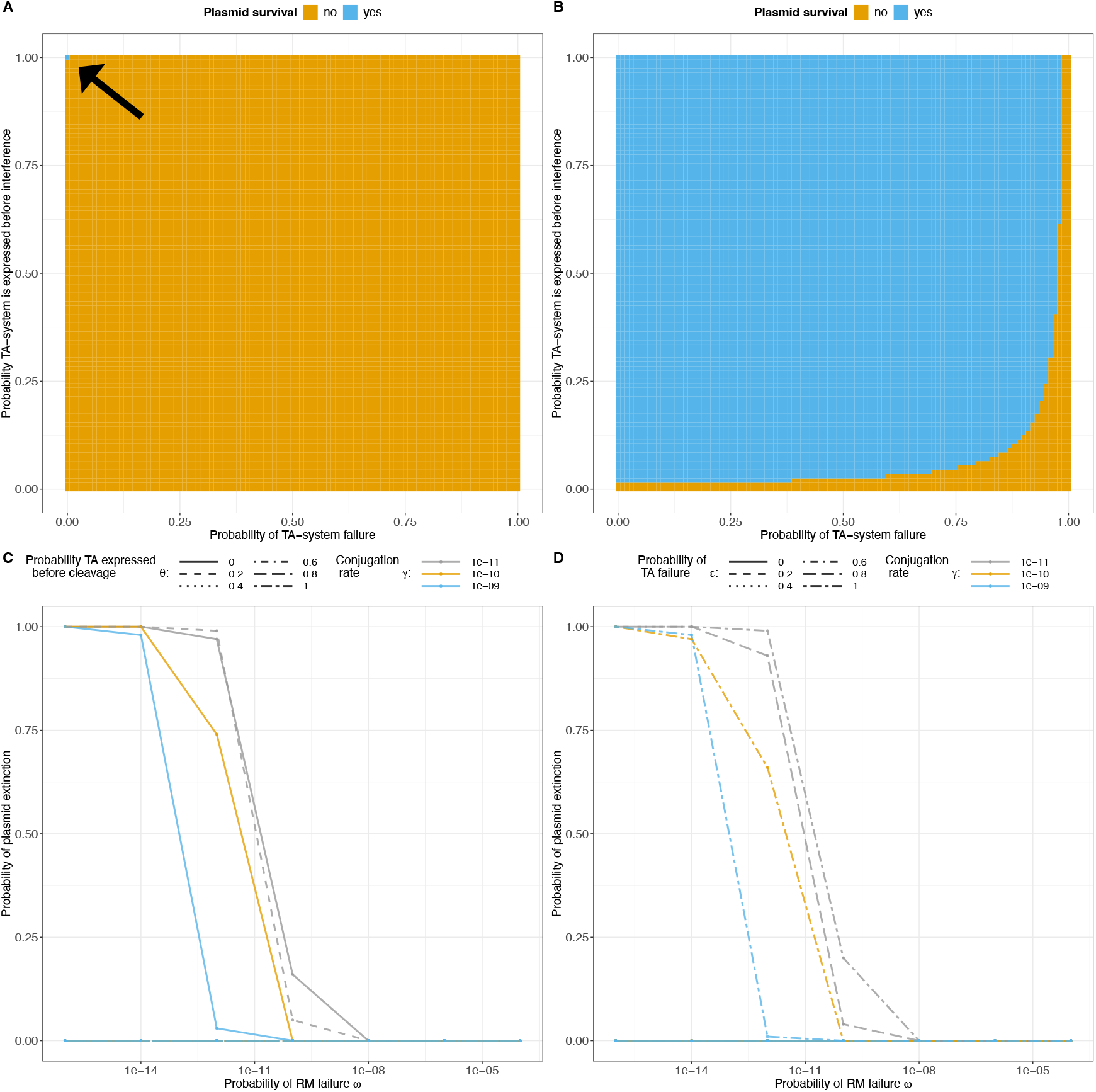
Impact of TA-systems on plasmid survival. **A, B**. The expression of TA-system before CRISPR-Cas interference favours plasmid survival. However, the mechanism of killing resistant cells through conjugation is more robust to TA-system failure and expression timing (**B**) than preventing the emergence of resistant cells (**A**). **C, D**. The expression of TA-system before RM interference favours plasmid survival. **C**. Plasmid extinction depends on the probability that the TA system is expressed before RM cleavage, represented by *θ*. The probability of TA failure is set to *ϵ* = 0. **D**. Plasmid extinction is relatively robust to the probability of TA failure, denoted as *ϵ*. We set the probability that the TA system is expressed before RM cleavage to *θ* = 1. In all simulations, we set the probability of losing a plasmid to *τ* = 10^−4^, and the plasmid fitness effect to *c* = 0.1 and in the CRISPR-Cas simulations the conjugation rate is set to *γ* = 10^−10^.

While no experimental evidence, to our knowledge, demonstrates the successful expression of plasmid-encoded TA systems before RM cleavage, resulting in cell death, the experimental setup suggesting TA expression before CRISPR-Cas interference raises parallels with innate bacterial immunity. During the experiments bacteria already carried a spacer targeting the plasmid prior to infection and consequently did not need to acquire a spacer [39]. Therefore, if RM cleavage kinetics are slower than the expression of plasmid encoded toxins — plausible given the presence of TA systems in the leading region of plasmids [25] — TA systems may facilitate plasmids in evading innate immunity. Expanding our basic RM-model to include TA-system dynamics (SI section 1.2) reveals that the expression of TA-system before plasmid degradation can also enhance the likelihood of plasmid survival in the context of RM-systems (figure 5C,D). This is because the conjugation of costly plasmids into cells, where the plasmid lacks the corresponding methylation pattern to be recognized as ‘self’, results in death of those cells. This decreases the probability of plasmid-carrying cells being outcompeted by fitter plasmid-free cells.

#### Bet/Exo

A recent discovery indicates that plasmids can evade CRISPR-Cas using the *Bet/Exo* system. [23] This system repairs double-stranded breaks caused by CRISPR-Cas interference through recombination. The plasmid sequence around the break is remodeled, allowing the plasmid to escape the spacer causing the break (for more detail, see SI section 2.4). A simplified mathematical model of *Bet/Exo*, considering a single CRISPR-Cas spacer (SI section 2.4), illustrates that *Bet/Exo* can allow the persistence of costly plasmids in the population (figure 6A,B). However, our model shows that costly plasmids can be maintained only if *Bet/Exo*-induced modifications do not substantially reduce the conjugation rate (figure 6C). Global sensitivity analysis demonstrates that the final plasmid prevalence in the population mainly depends on the effect of the modification on conjugation rate followed by the plasmid’s effect on bacterial growth and segregation loss (figure 6D, SI section 2.4.2). The probability with which *Bet/Exo* successfully modifies the plasmid before degradation has only a weak effect on the plasmid frequency in the long run (figure 6D). This suggests that if the modification of *Bet/Exo* allows plasmid maintenance, the dynamics are primarily driven by the presence of the modified plasmid rather than the frequency with which *Bet/Exo* modifies the plasmid.

**Figure 6.**
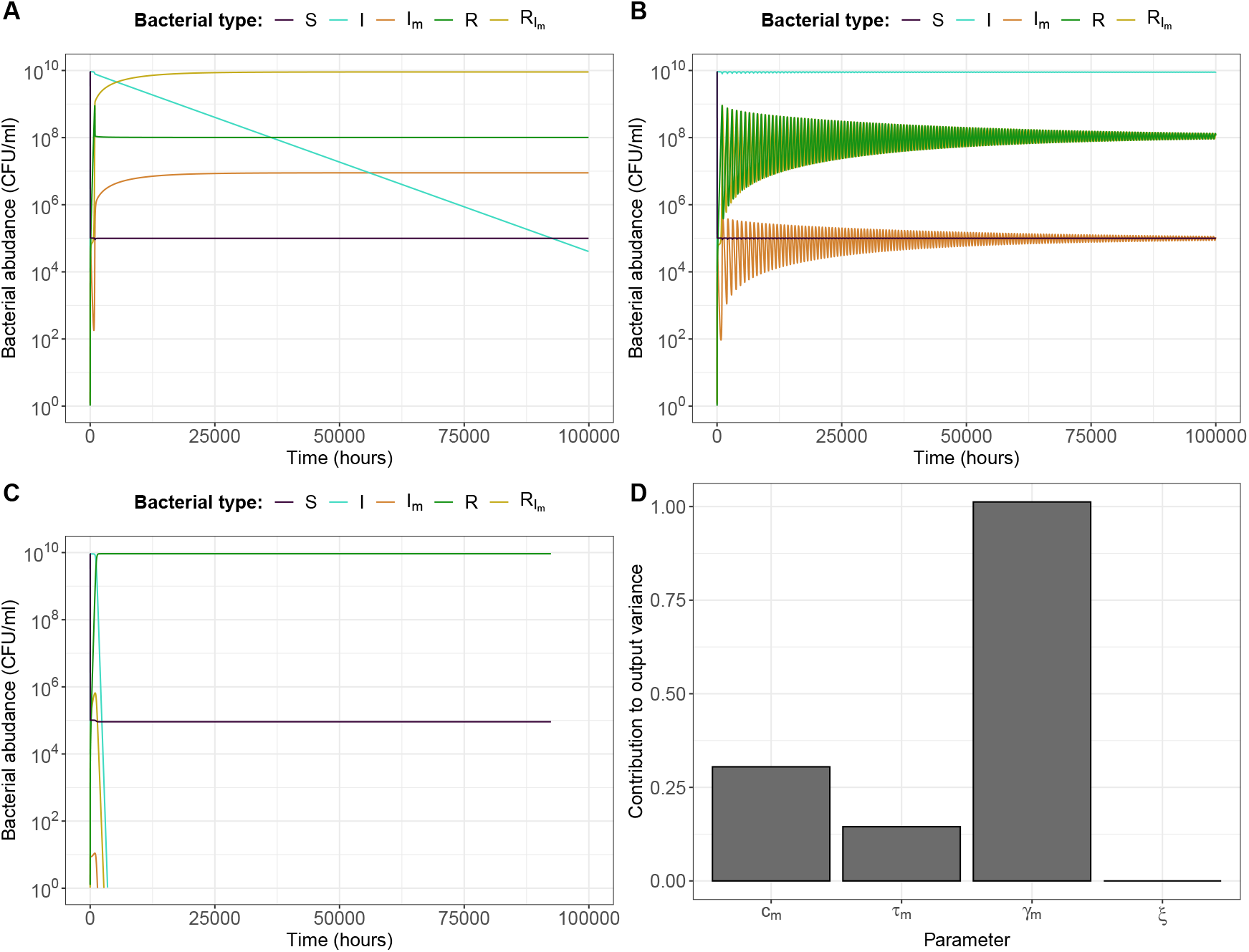
Impact of *Bet/Exo* plasmid modifications on plasmid prevalence. Scenarios (**A**) and (**B**) illustrate two exemplary outcomes where *Bet/Exo* modifications lead to the maintenance of costly plasmids: **A**. The modified plasmid is able to dominate in the bacterial population despite higher segregation loss as it has the advantage of infecting resistant cells. **B**. The population is dominated by the wildtype plasmid as the *Bet/Exo* modification affects the plasmid’s effect on bacterial growth, disadvantaging the resistant population infected with the modified plasmid in growth competition. **C**. The plasmid goes extinct due to the *Bet/Exo* modification, which reduces the conjugation rate to an insufficient level. The bacterial population consists of CRISPR-Cas cells naïve to the plasmid *S*, naive cells infected with the wild-type plasmid *I*, naive cells infected with the modified plasmid *I*_*m*_, cells resistant to the wild-type plasmid *R*, and resistant cells infected with the modified plasmid 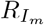. In the simulations, we set the conjugation rate to *γ* = 10^−10^, the probability of losing a plasmid to *τ* = 10^−4^, the plasmid fitness effect to *c* = 0.1, and the *Bet/Exo* modification probability before CRISPR-Cas interference to *ξ* = 0.8. The *Bet/Exo* modification affects only one plasmid trait at a time, with others remaining identical to the wild-type. In (**A**), *τ*_*m*_ = 10^−1^ for the probability of losing the modified plasmid; in (**B**), *c*_*m*_ = 0.2 for modified plasmid fitness effect; and in (**C**), *γ* = 10^−13^ for modified plasmid conjugation rate. **D**. Total order Sobel indices indicating the importance of parameters to explain plasmid prevalence. *c*_*m*_ is the modified plasmid fitness effect, *τ*_*m*_ the probability of losing the modified plasmid, *γ* the modified conjugation rate, *ξ* the probability of *Bet/Exo* modification before CRISPR-Cas interference. Note that total order Sobol indices also includes interactions of parameters and therefore does not necessarily add up to one. See SI section 2.4.2 for a detailed explanation and parameter bounds used for the global sensitivity analysis.

If the plasmid is maintained in the population, the impact of the modification on plasmid traits also determines the prevalence of the modified plasmid versus the wild-type plasmid in the long run. The modified plasmid can dominate if *Bet/Exo* modifications lead to a decrease in segregation loss or conjugation rate. In this case, the modified plasmid retains a selective advantage, as it can still infect resistant cells (figure 6A). The wild-type plasmid may dominate if the modified plasmid imposes a higher cost on bacterial growth (figure 6B). Despite the modified plasmid’s ability to infect resistant cells, those resistant cells experience a growth disadvantage compared to cells infected with the wild-type plasmid, keeping the modified plasmid and cells resistant to the wild-type plasmid at low frequencies.

#### Evasion of CRISPR-Cas spacers through plasmid evolution

It is important to note that in our basic CRISPR-Cas model we make a strong assumption that the plasmid does not evolve, despite the two well-known facts: (i) plasmids can mutate and undergo evolutionary change [40],and (ii) a single point mutation in the protospacer is enough to escape a spacer [41]. To explore the impact of plasmid evolution on plasmid survival in the presence of CRISPR-Cas, we developed a mathematical model that considers some complexities in the dynamics of CRISPR-Cas and plasmids. Our model includes population-wide spacer heterogeneity, allowing each cell to acquire one plasmid spacer, as population-wide spacer diversity has been shown to be key for the efficiency of CRISPR-Cas against virulent phages [6]. We also consider the potential for plasmids to escape spacers through mutation (further details in SI section 2.3). We find that population-wide spacer heterogeneity, which is often sufficient to lead phages to extinction within a few days [6, 42], can substantially lower plasmid prevalence without driving plasmids to extinction (figure 4C). The reasons for this observation require further investigation, and it is important to note that our current findings are based on a limited parameter set. Within the explored parameter space, our results suggest that, although the plasmid persists, the bacterial population benefits from CRISPR-Cas immunity through a substantial reduction in plasmid prevalence.

## 4 Discussion

In our study, we investigate the dynamics of plasmid spread in the presence of bacterial innate immunity (RM-systems) and adaptive immunity (CRISPR-Cas). Our results show that beneficial plasmids can persist in a bacterial population despite the challenges posed by bacterial immunity. However, for costly plasmids, we find that bacterial immunity can substantially reduce their prevalence and may even lead to their extinction. The efficiency of bacterial immunity against plasmids can be strongly determined both by the strength of immunity and the plasmid’s fitness. We further illustrate how escape mechanisms, such as TA-systems, the *Bet/Exo* system, and evasion of CRISPR-Cas spacers through plasmid evolution can mitigate the burden imposed by bacterial immunity on plasmids.

Our finding that beneficial plasmids can persist in the presence of bacterial immunity is supported by a recent study showing that, under antibiotic pressure, resistance plasmids can persist in the presence of CRISPR-Cas [43]. Additionally, our result, suggesting that bacterial immunity can impose a significant burden on plasmids, aligns with previous bioinformatic studies and experimental findings [15, 20, 43]. Plasmids often carry genes that enable evasion of bacterial immunity [20, 21, 25], reinforcing the idea that these defense systems can be challenging for plasmids. However, our results suggest that the effectiveness of bacterial immune systems in blocking plasmids strongly depends on whether these plasmids confer a benefit or cost to bacterial cells. Previous work based on CRISPR-Cas suggests that in the presence of beneficial plasmids, the loss of immunity function might be favored [12, 29]. However, losing the function of bacterial immunity could be a major disadvantage in natural environments, where bacteria are likely to frequently interact with phages due to their high abundance [30]. Our results demonstrate that, even in environments with selection pressures to maintain functioning bacterial immunity, beneficial plasmids can maintain in a bacterial population.

While our results generally indicate similar qualitative outcomes for plasmid prevalence in the presence of RM or CRISPR-Cas, the dynamics of plasmid spread differ. Depending on the benefit or cost conferred by the plasmid on bacteria, RM and CRISPR-Cas provide distinct advantages or disadvantages, due to their innate and adaptive nature. RM-systems are immediately effective; however, if the costly plasmid can overcome the barrier of RM-systems, it faces no further burden. In contrast, with CRISPR-Cas, the plasmid can initially spread in the bacterial population irrespective of its fitness effect on the bacteria. Although it may take time for CRISPR-Cas to significantly decrease plasmid prevalence, its adaptive ability to acquire different spacers imposes a constant pressure on plasmids. However, in the case of a beneficial plasmid, this adaptive ability results in a growth disadvantage for the bacterial population, as it produces less fit cells resistant to the plasmid. The immediate barrier imposed by RM-systems can also lead to a disadvantage in the case of beneficial plasmids. Low probabilities of RM failure in bacterial strains, reflecting strong RM systems, can reduce bacterial population diversity and, consequently, RM diversity. This reduced RM diversity may disadvantage the population in future infections with harmful genetic elements, especially phages [2,44].

Several mechanisms allow costly plasmids to lower the burden of bacterial immune systems (table 1). We show that expressing TA-systems before immune interference ensures plasmid maintenance in the presence of RM and CRISPR-Cas, aligning with Van Houte et al.’s hypothesis that this mechanism explains plasmid survival despite CRISPR-Cas [39]. However, in case of RM system it favours decreased bacterial diversity, which may be a disadvantage for future infections of costly mobile genetic elements [2, 44]. The overall importance of TA-systems in evading bacterial immunity is yet to be fully assessed, as chromosomes also frequently carry TA-systems. If the cell encodes a suitable anti-toxin on the chromosome, it could serve as an ‘anti-addiction’ module [45] and compromise the effectiveness of TA-systems as an escape mechanism.

Our results illustrate that *Bet/Exo* can be an effective mechanisms for costly plamsids to evade CRISPR-Cas immunity. However, the effectiveness of the *Bet/Exo* mechanism in maintaining costly plasmids is closely tied to its impact on the plasmid’s conjugation rate (figure 6C,D). The reduction in conjugation rate resulting from *Bet/Exo* modifications is likely influenced by whether the spacer targets essential genes for conjugation, as supported by experimental data [23]. Consequently, the maintenance of costly plasmids by *Bet/Exo* relies on chance — specifically which spacer CRISPR-Cas acquires. Especially in the long-term, when CRISPR-Cas can acquire multiple spacers, it is likely that a spacer is acquired where *Bet/Exo* modifications do not lead to plasmid maintenance. Whether *Bet/Exo* confers also an advantage for overcoming the barrier imposed by RM-systems remains unclear.

By repairing double-stranded breaks caused by RM-systems, *Bet/Exo* could increase the failure rate of RM-systems, making it more likely that plasmids get fully methylated and recognized as ‘self’ before being degraded. This could be an effective strategy to evade RM systems, especially for smaller plasmids, which are likely to experience fewer double-stranded breaks due to RM interference compared to larger plasmids [20]. If *Bet/Exo* indeed leads to an increased failure rate of RM-systems, it could help plasmids in overcoming the barrier imposed by RM-systems, depending on the plasmid fitness effects of *Bet/Exo* modification (figure 3). However, the impact of *Bet/Exo* on plasmid spread and survival in the presence of RM-systems needs to be experimentally tested.

Plasmid evolution can lead to evasion of bacterial immunity through mutations in three distinct ways: (i) compensation of plasmid cost, (ii) increase of conjugation rate, (iii) escape of immune targets. First, our results show that the elimination of plasmids is closely tied to the plasmid fitness cost, with low-cost plasmids taking weeks to be eradicated (figure 4A, figure S1A). Laboratory experiments have shown that compensatory mutations increasing the fitness of plasmid-carrying cells can be observed within a few days [46–49]. Some of these mutations fully compensate for the plasmid cost or even render plasmid-carrying cells fitter than the plasmid-free wild-type cell [47–49]. Consequently, plasmids can be maintained if their fitness cost is eliminated by compensatory mutations before being out-competed. It is important to note, that compensatory mutations ameliorating plasmid cost can occur not only on the plasmid itself but also on the chromosome [50].

Second, our results show that, for RM-systems, mutations leading to a higher conjugation rate can enhance the probability of plasmid survival, depending on the probability of RM failure (figure 3B). However, such an increase in the conjugation rate typically comes with a trade-off in plasmid fitness cost [51]. Our findings show that a higher plasmid cost increases the probability of plasmid extinction (figure 3A). Therefore, whether an increase in conjugation rate ensures plasmid survival depends on how much costlier the plasmid becomes for the bacterial host through the increase of conjugation rate. In the case of CRISPR-Cas, our basic model suggests that an increase in the conjugation rate does not confer a benefit in overcoming the immunity barrier.

Third, our model taking into account the co-evolution between CRISPR-Cas and plasmids, shows that plasmid mutations, enabling the escape from CRISPR-Cas spacers, can ensure plasmid maintenance. Our investigated parameter-set and our model only takes into account some part of the complexity between CRISPR-Cas and plasmids co-evolution. Therefore, the robustness of our result, indicating that costly plasmids can persist, even if just at a low frequency, requires further investigation. We make three main simplifications in the simulations of the co-evolution between plasmids and CRISPR-Cas, and considering these could potentially alter predictions regarding plasmid maintenance. First, our model assumes a maximum of one spacer per cell, which can be sufficient to lead phages to extinction [6, 42]. As within-cell spacer-heterogeneity is even more efficient than population wide-spacer heterogeneity [5], considering multiple spacers per cell could potentially lead plasmids to extinction. Second, while plasmids typically have multiple copies within a cell, we assume these copies are identical. In reality, heterogeneity between plasmid copies due to mutations exists [40, 52], potentially influencing spacer effectiveness. However, our understanding of the dynamics of plasmid mutations and heterogeneity over time remains limited [52]. Third, we assume that all mutations negatively affect plasmid growth, yet the location of mutations in the plasmid genome may vary in terms of their impact on fitness.

Mutations in essential genes are more likely to affect plasmid survival, similarly as for *Bet/Exo*. Studies show that spacer targets are not randomly distributed among the plasmid genome [53, 54]; rather, they often specifically target essential plasmid backbone genes, such as those involved in replication and conjugation [53]. If we were to consider varying fitness effects of plasmid mutations, including the possibility to affect the conjugation rate, a costly plasmid might face extinction when CRISPR-Cas acquires a spacer that is challenging to evade through mutation without causing a substantial impact on plasmid fitness. This could occur, for example, if the mutation substantially reduces plasmid conjugation, as demonstrated in the case of *Bet/Exo*. Overall, the degree to which plasmid evolution, enabling evasion of CRISPR-Cas, serves as a survival mechanism remains uncertain. Our understanding of CRISPR-Cas and plasmid interactions, especially co-evolutionary dynamics, and the differences compared to phages, is limited. In the context of RM-systems, plasmid evolution is an unlikely strategy to overcome the barrier imposed, given that conjugative plasmids typically carry multiple RM restriction targets [20].

For both RM-Systems and CRISPR-Cas, specific proteins known as anti-RM and anti-CRISPR proteins, respectively, are known to suppress the function of these immune systems [22, 55]. Indeed, experimental data demonstrate that anti-CRISPR proteins can protect plasmids from CRISPR-Cas targeting [21]. Additionally, plasmid-encoded anti-RM systems have been shown to compromise the efficiency of RM systems [56] and lower the dissemination of plasmids [57, 58]. In case of RM system the effectiveness of anti-RM can easily be illustrated with our basic RM-model by a high RM-failure probability (figure 3). However, a major disadvantage of this evasion strategy is that not only the plasmid itself could benefit from the suppressed immunity but also other competing plasmids or virulent phages [59–61], which eventually could lead the plasmid to extinction.

Overall, our study highlights, that RM and CRISPR-Cas are indeed a barrier for costly plasmids, imposing evolutionary pressure on them. However, the dynamics between bacterial immunity and plasmids can be complex, enabling costly plasmids to endure in the population for a period before facing potential exclusion. The pressure of bacterial immunity for costly plasmids can be mitigated by several escape mechanisms. Importantly, we demonstrate that despite bacterial immunity, beneficial plasmids — key drivers of bacterial evolution — can still be maintained. Nevertheless, our understanding of the interactions of bacterial immunity and plasmids is in its early stages. Our basic mathematical framework provides a foundation for future exploration and refinement to enhance our comprehension of these dynamics. For instance, our basic RM model does not take into account all the complexity of a bacterial population consisting of different RM systems. Bacterial types may share common RM systems [7] or have sets of RM-systems with different failure probabilities, and plasmid traits, such as conjugation, segregation loss and the effect on growth, could be influenced by the type of bacteria [50]. Considering these factors could enhance our understanding of the dynamics between plasmids and bacterial defence mechanisms and some of those factors are expected to provide further explanations why bacteria commonly carry costly plasmids despite bacterial immunity. For instance, the varying plasmid fitness effects among different hosts within a bacterial community can enhance plasmid persistence [62] and may contribute in overcoming the barrier posed by RM-systems. Bacterial hosts to which the plasmid is well adapted can act as a constant plasmid source, providing continuous opportunities for plasmids to overcome the RM-system barriers of other bacterial hosts. Furthermore,in this work, we hypothesize that phage pressure prevents the loss of innate or adaptive immunity. Explicitly modeling phage dynamics and immune loss, would allow to study how strong and frequent phage pressure needs to be to prevent immune loss, despite the potential cost of a small bacterial subpopulation targeting beneficial plasmids. Finally, although our work does not address the potential synergy of CRISPR-Cas and RM systems within a single bacterial cell [63, 64], experimental findings suggest that this interplay enhances the barrier against conjugative plasmids [15]. Extending our proposed models to incorporate the combined effect of bacterial innate and adaptive immunity may provide valuable insights and allow to assess whether this co-occurrence provides a more robust barrier against plasmids.

## Data availability

Data from stochastic simulations and code are available available at OSF: https://osf.io/r6uzd/?view_only=a4b6fc8348e748e6ae381da564d96873.

## Acknowledgments

We thank Eduardo Rocha for helpful comments on the manuscript. We thank Christopher Witzany for helpful discussion and providing references. We also thank Sneha Sundar, Ricardo Leon Sampedro and the theoretical biology group for helpful input. We used the assistance of ChatGPT 3.5 for checking and refining our writing as well as for advice on plotting aesthetics and code debugging. We used Perplexity AI to search for relevant references.

## Competing Interests

The authors declare no competing interests.

## Notes

https://osf.io/r6uzd/?view_only=a4b6fc8348e748e6ae381da564d96873

### Competing Interest Statement

The authors have declared no competing interest.

