## Supplementary materials for "Assessing the role of bacterial innate and adaptive immunity as barriers to conjugative plasmids"

Berit Siedentop[1*], Carlota Losa Mediavilla[1], Roger D. Kouyos[2,3], Sebastian Bonhoeffer[1†], Hélène Chabas[1†]

**1** Institute for Integrative Biology, ETH Zürich, Zürich, Switzerland
**2** Division of Infectious Diseases and Hospital Epidemiology, University Hospital Zurich, University of Zurich, Zurich, Switzerland
**3** Institute of Medical Virology, University of Zurich, Zurich, Switzerland

†These authors contributed equally

*

### 1 Restriction-modification systems

#### 1.1 Basic model

##### 1.1.1 Model description

**Bacterial population composition** We consider a population of bacterial strains that uses Restriction-modification (RM) systems as a defence mechanism against foreign DNA and compete for the same resource. We assume that within the bacterial population there exists a heterogeneity of $n$ sets of different RM-systems. A bacterial cell that carries the set of RM system $i$ is denoted by $b_i$, with $i \in \{1, \ldots, n\}$. We additionally consider a conjugative plasmid in the bacterial population that is able to infect a bacterial cell with RM system $i$ in case the plasmid has the corresponding methylation pattern. We only consider one plasmid variant, which can only differ in its mythelation pattern. A bacterial cell with a set of RM-Systems $i$ that is infected with a plasmid with the corresponding methylation pattern is denoted by $b_i^I$.

**Bacterial population size** The bacterial growth is modeled by a density dependent division rate $\rho_i \left(1 - \frac{B+B^I}{K}\right)$, where $\rho_i$ is the maximal division rate of bacterial cells $b_i$, $K$ the carrying capacity, and $B = \sum_{i=1}^{n} b_i$ and $B^I = \sum_{i=1}^{n} b_i^I$ the total bacterial cell density of plasmid free and plasmid carrying cells respectively. Plasmids can not only carry beneficial accessory genes that encode resistances against antibiotics or heavy metals, but it has also been reported that plasmids encode genes, which allow the usage of new energy resources such as lactose, other sugar derivatives or light energy [1–3]. We assume that the conjugative plasmid can enable the use of a new energy source, which is reflected in our model with a higher carrying capacity $K_I$ for bacterial cells that carry the plasmid than for cells that do not carry a plasmid ($K_I > K$). Plasmid carrying cells can use the same resources as plasmid free cells and additionally another resource. The factor $\chi$ reflects how much plasmid free cells $b_i$ participate in the resources of plasmid carrying cells $b_i^I$. The density dependent growth rate for plasmid carrying cells is therefore described by: $\rho_i\,(1-c)\left(1 - \frac{B_I+\chi B}{K_I}\right)$, where $c$ denotes the fitness effect of a plasmid ($c > 0$: cost, $c < 0$: benefit). We assume that bacterial cells $b_i$ die with a constant rate $d_i$. With the density dependent growth rate and constant death rate the bacterial population size is bounded, however once this bound is reached the plasmid fitness effect can still have an effect on the frequency of individual cell types.

**Segregation loss and conjugation** Cell division of plasmid carrying cells $b_i^I$ may not necessary lead to two plasmid carrying daughter cells. At cell division the plasmid gets lost from one daughter cell with probability $\tau$, which is called segregation loss. We assume a well mixed bacterial population, where plasmid conjugation occurs between plasmid free cells $b_i$ and plasmid carrying $b_i^I$ in a density-dependent manner. As the bacterial population consists of $n$ different types of bacteria with unique RM-systems the identity of donor and recipient also determine the rate of transmission. In case the donor and recipient carry the same set of RM-Systems $i$ the conjugation rate is $\gamma$ as the plasmid already has the corresponding methylation pattern. However, in case donor and recipient carry different types of RM-Systems, the RM-System of the recipient is a barrier to successful plasmid conjugation. The incoming plasmid will have a methylation pattern that is recognized as foreign and therefore the plasmid will be degraded. But RM-systems have a certain failure probability $\omega_i$, where the incoming plasmid will be methylated to the new host's metyhlation pattern, before the plasmid is cleaved. Conjugation between bacterial cells of different types of RM-systems occurs therefore rarely with the rate $\omega_i\gamma$. Note that we neglect that conjugation rate between different bacterial types could also differ due to other reasons than RM-systems. We also neglect that the plasmid segregation loss, and plasmid fitness effect on bacterial growth could vary between different cell types. [4]

The described processes can be depicted in a schematic flow diagram (main text figure 1A,C) and captured by the following equations:

$$
\begin{aligned}
\frac{\mathrm{d}b^i}{\mathrm{d}t} =& \rho_i b^i \left(1-\frac{B+B^I}{K}\right) + \tau_i\rho_i\left(1-c_i\right) b_i \left(1-\frac{B_I+\chi B}{K_I}\right) - \left(\sum_{\substack{j=1\\ j\neq i}}^{n} \gamma_{j,i}\omega_i b_I^j + \gamma_i b_I^i\right) b^i - d_i b_i, \\
\frac{\mathrm{d}b_I^i}{\mathrm{d}t} =& \rho_i\left(1-c_i\right)\left(1-\tau_i\right) b_I^i \left(1-\frac{B_I+\chi B}{K_I}\right) + \left(\sum_{\substack{j=1\\ j\neq i}}^{n} \gamma\omega_i b_I^j + \gamma_i b_I^i\right) b^i - d_i b_I^i.
\end{aligned}
\tag{1}
$$

##### 1.1.2 Important model assumption about RM-systems

For phages, the probability of RM escape has been estimated to range from $10^{-2}$ to $10^{-8}$ [5,6]. To our knowledge, there is currently no estimate for plasmids. Bacterial cells are often found to carry multiple distinct RM systems per cell [6,7]. In our model, multiple distinct RM-systems per cell can be represented with a lower RM failure probability $\omega_i$. When a cell carries multiple RM-systems it often carries two distinct RM-system [6]. Assuming that the failure of those systems is independent, the total probability for RM failure is the product of the individual failure probabilities. Considering $10^{-8}$ as the lowest failure probability for one RM system, we obtain $10^{-16}$ as the lowest failure probability for two RM systems, serving as the minimum RM failure probability in our simulations.

Besides the possibility of within host heterogeneity of RM systems it is known that a bacterial population can also show a population wide RM heterogeneity of distinct RM systems [6,7]. Often bacterial strains do not have any common RM-systems [6], we therefore simplified our model and exclude that strains could partially carry overlapping RM-systems.

As a simplification for our simulations, we further assume, even though each bacterial cell carries distinct RM-systems, that the probability of RM failure for each bacterial species $\omega_i$ is the same.

Note that the potential cost of maintaining RM-systems is incorporated in our model through the maximal cell division rate. We assume that all bacterial cells in the population carry RM-systems and assume that distinct RM-systems confer the same cost to the bacterial cell.

##### 1.1.3 Implementation

For RM simulations, we stochastically implement the set of equations 1. In the context of RM-systems, stochastic and deterministic implementations can result in qualitative different outcomes, especially for costly plasmids (figure S1). In the stochastic implementation, conjugation events that may lead to overcoming the RM-system barrier occur stochastically, introducing a potentially longer waiting period before a plasmid can enter a new bacterial species and acquire the corresponding methylation pattern. During this time, plasmid-carrying cells face the risk of being outcompeted by plasmid-free cells. Moreover, in the stochastic implementation, even if a plasmid successfully overcomes a barrier and acquires the necessary methylation in a new bacterial strain, it might still be lost in this strain due to extinction by chance.

In all our simulations, we make the following assumptions: each bacterial species has a maximal cell division rate of $\rho_i = 1.3$, and a death rate of $d_i = 0.1$. The carrying capacity is set to $K = 10^{10}$. Additionally, unless explicitly stated otherwise, we assume that the plasmid does not confer the ability to utilize an additional resource, meaning $K = K_I$ and $\chi = 1$. We initialize our simulations at the maximal population size, where one of the bacterial strains carries the plasmid in small concentration $b_1^I = 10^4 \frac{\text{CFU}}{\text{ml}}$, and the rest of the population size is equally distributed between the plasmid-free carrying strains. The simulations are run for a maximum of $10^5$ hours, but are terminated earlier in case the plasmid goes to extinction or the plasmid is able to enter each individual bacterial strain, i.e., $b_i^I > 100 \frac{\text{CFU}}{\text{ml}}$ for each $i$. Each parameter set we simulate a 100 times based on which we calculate the probability of plasmid extinction.

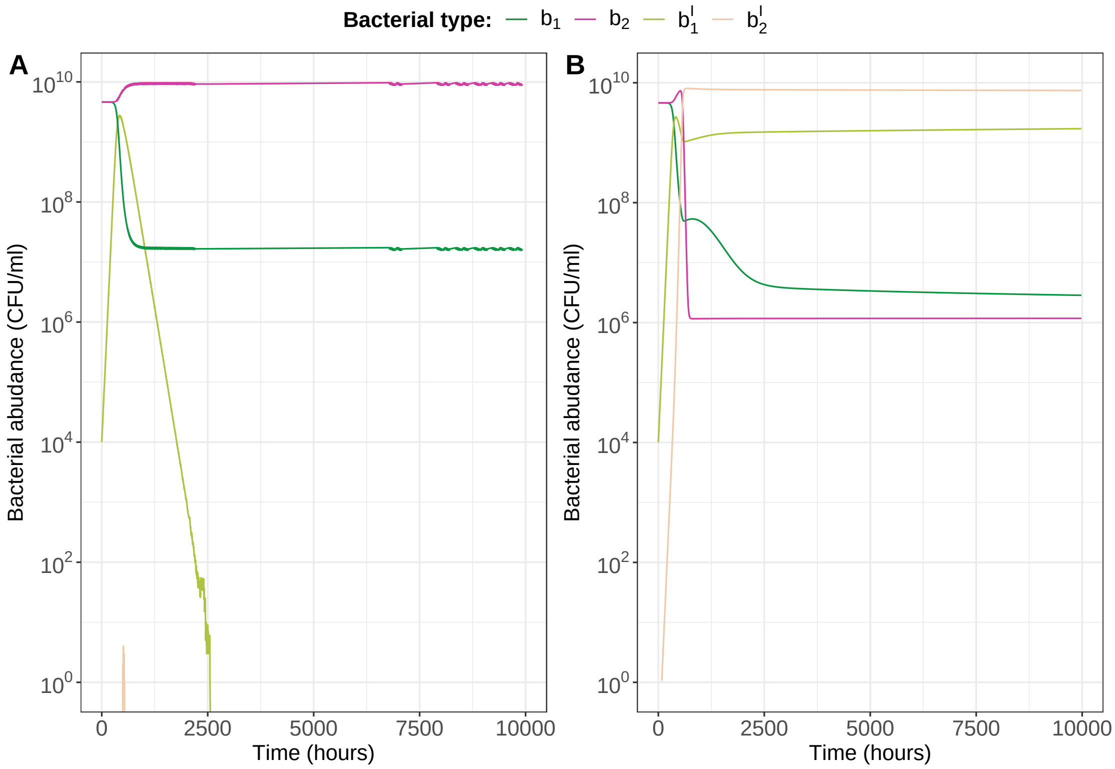


**Figure S1. Stochastic and deterministic implementation of the RM-model can lead to different qualitative behaviour.** (**A**) and (**B**) are stochastic and deterministic implementations of the RM-Model with the same set of parameters, where in the stochastic simulation (**A**) the plasmid goes to extinction, whereas in the case of the deterministic simulation (**B**) the plasmid is maintained in the population. In the simulations, we set the conjugation rate to $\gamma = 10^{-11}$, the probability of losing a plasmid to $\tau = 10^{-4}$, the plasmid fitness effect to $c = 0.1$, and the probability of RM-failure to $\omega = 10^{-11}$.

##### 1.1.4 The maintenance of beneficial plasmids is robust to variations in plasmid traits and strength of RM immunity

Our simulations show that the maintenance of plasmids with positive effect on bacterial growth is robust to variations in plasmid traits, i.e. conjugation rate and segregation loss, and the strength of RM immunity (figure S2). The strength of RM immunity is determined in our model by the probability of RM failure $\omega$.

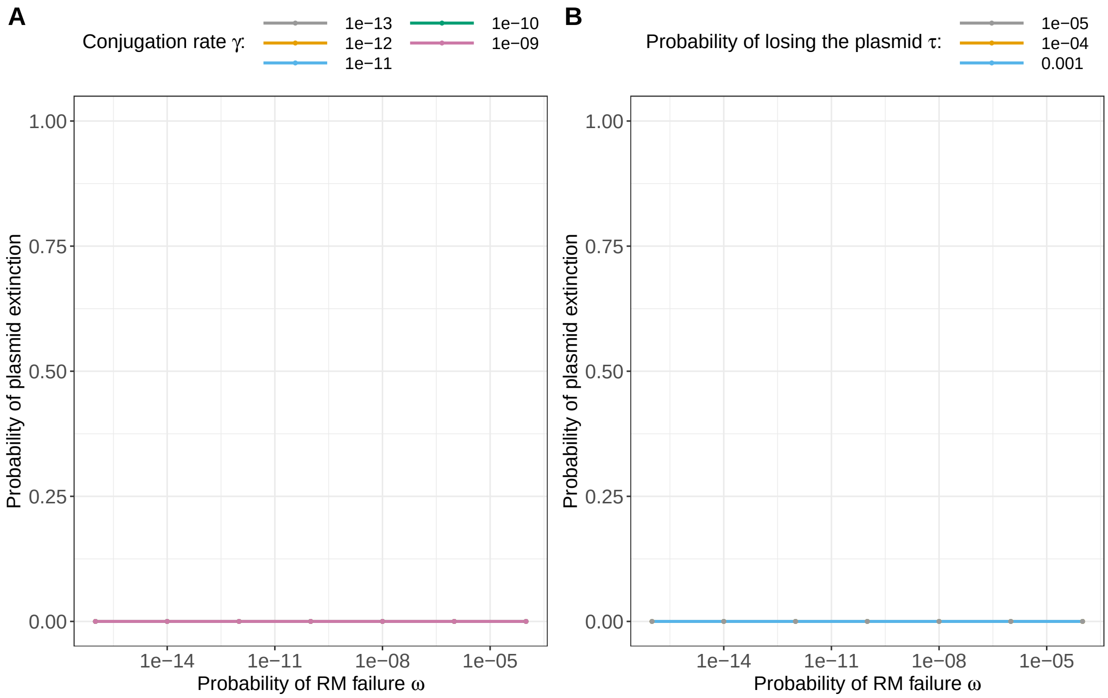


**Figure S2. The maintenance of plasmids with positive effect on bacterial growth is robust to variations in plasmid traits and strength of RM immunity. A.** Probability of plasmid extinction in dependence on conjugation rate and RM failure. **B.** Probability of plasmid extinction in dependence on segregation loss and RM failure. Plasmid fitness effect is set to $c = -0.05$ in all simulations. For panel (**A**), the probability of losing the plasmid is $\tau = 10^{-4}$, and for panel (**B**), the conjugation rate is $\gamma = 10^{-11}$.

##### 1.1.5 Beneficial plasmids can alter strain diversity

Our results show that beneficial plasmids can be maintained in a bacterial population in the presence of RM-systems. There are two scenarios: (i) the plasmid spreads to other bacterial strains (figure 2A), or (ii) the plasmid-carrying strain out-competes others due to its failure to overcome the barriers imposed by RM-systems in time (figure 2B). The likelihood of each scenario occurring depends on the plasmid's ability to overcome individual barriers of distinct RM-systems in plasmid-free bacteria and establish itself in that population before being out-competed by already plasmid-carrying bacteria. Scenario (i) is favored with higher conjugation rates, higher RM failure probability, and a lower increase in the maximal cell division rate due to the plasmid, wheres scenario (ii) is favored with lower conjugation rates, higher RM failure probability, and a higher increase in the maximal cell division rate due to the plasmid (figure S3). Higher conjugation rates provide more opportunities for the plasmid to overcome RM-system barriers, and lower RM failure probability increases the likelihood of successful plasmid entry during conjugation events. Additionally, a lower increase in the maximal cell division rate caused by the plasmid extends the time for the plasmid-carrying strain to out-compete plasmid-free strains, thereby increasing the window of opportunity for plasmid spread.

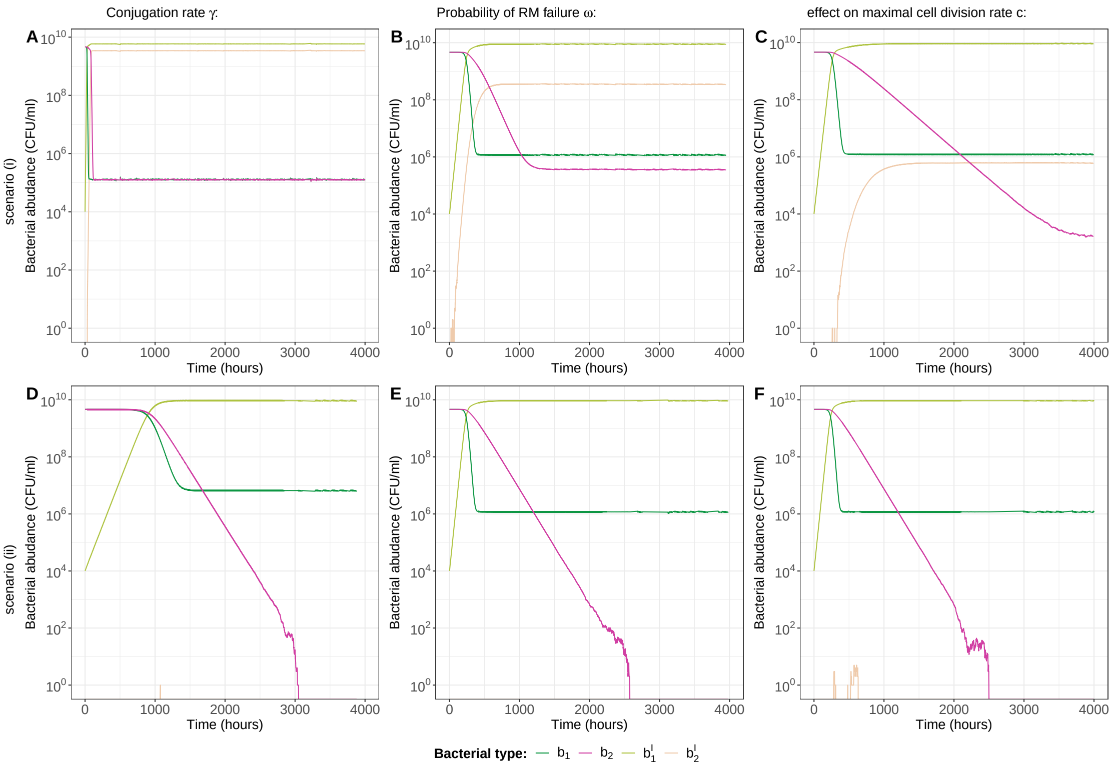


**Figure S3. The fitness of beneficial plasmids and the strength of immunity can influence strain diversity in the bacterial population.** The three columns illustrate the impact of conjugation rate, probability of RM failure, and the plasmid effect on the maximal cell division rate on bacterial strain diversity resulting from the spread of a beneficial plasmid. The plasmid spread to other bacterial strains, and therewith strain diversity, is favoured with higher conjugation rates (**A**), higher probabilities of RM failure (**B**) and lower benefit for cell division (**C**). Conversely, low strain diversity is favoured with lower conjugation rates (**D**), lower probabilities of RM failure (**E**) and higher benefit for cell division (**F**). Plasmid fitness effect is set to $c = -0.1$ except in panel (**C**), where it is set to $c = -0.05$. Plasmid conjugation rate is set to $\gamma = 10^{-11}$ except in panels (**B**) and (**E**), where it is set to $\gamma = 10^{-10}$ and $\gamma = 10^{-12}$, respectively. RM failure probabilities is set to $\omega = 10^{-10}$, except in panels (**B**) and (**E**), where they are set to $\omega = 10^{-5}$ and $\omega = 10^{-14}$, respectively.

##### 1.1.6 Plasmids allowing the utilization of an additional resource

When investigating the plasmid's ability to invade and persist in a bacterial population with RM protection, we observe that plasmids allowing the utilization of an additional resource can favour plasmid survival, even in circumstances where the plasmid would not have otherwise persisted. Specifically, we find that even when the barrier imposed by the RM system is high, as indicated by a low RM failure probability ($\omega = 10^{-10}$), a negative fitness effect of the plasmid on cell division ($c = 0.1$), and a conjugation rate of $\gamma = 10^{-11}$, the plasmid is able to persist (figure S4). This contrasts with our previous observation (main text figure 3A), where under these conditions, where the plasmid did not allow the utilization of an additional resource, we observed a probability of plasmid extinction of 100%. Despite the plasmid's impact on reducing maximal cell division, utilizing the additional resource provides an advantage, especially at high densities when the commonly used resource faces high competition among both plasmid-free and plasmid-carrying cells, leading to increased net-cell growth in plasmid-carrying cells.

Similar to plasmids that increase the maximal growth rate of bacteria, we observe that plasmids enabling the utilization of an additional resource can decrease strain/RM diversity (section 1.1.5, figure S4).

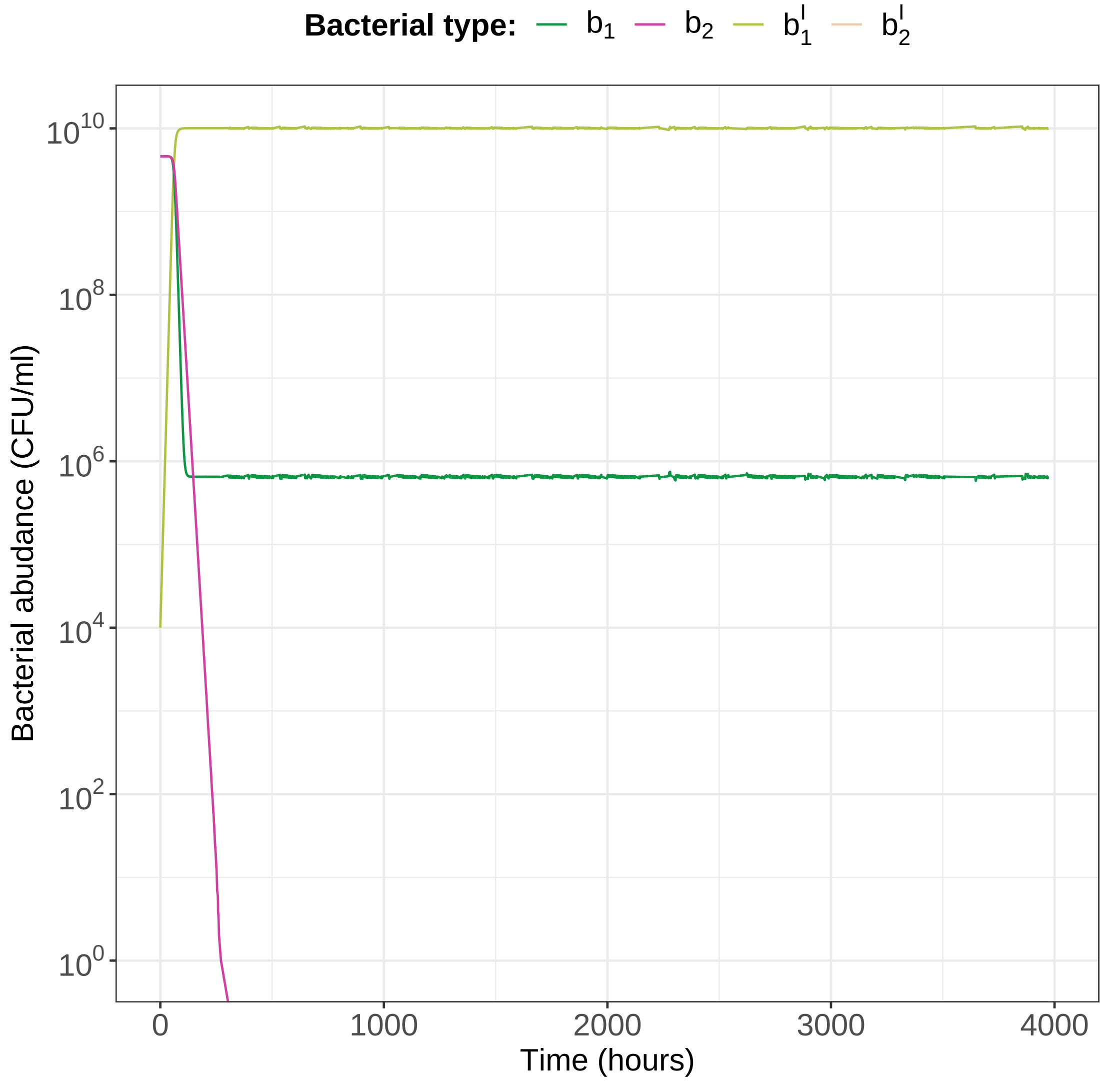


**Figure S4. Plasmids allowing the utilization of an additional resource favour plasmid maintenance** Exemplary simulation of a plasmid that allows the utilization of an additional resource. The carrying capacity of the plasmid carryin cells is increased by a factor of 1.1 (i.e., $K_I = 1.1$). The parameter $\chi$, indicating the extent of participation of plasmid-free cells in the resources of plasmid-carrying cells, is given by $\chi = \frac{K}{K_I}$. In our simulations, the plasmid has a negative effect on the maximal cell division rate ($c = 0.1$), the probability of RM failure is set to $\omega = 10^{-14}$, the conjugation rate is set to $\gamma = 10^{-11}$, and the probability of losing a plasmid to $\tau = 10^{-4}$.

#### 1.2 RM-model with TA system

**Bacterial population composition** As in the basic RM-model (section 1.1), the bacterial population consist of plasmid-free bacterial strains carrying the the unique set $i$ of RM-systems, with $i \in \{1, \ldots, n\}$. Bacterial cells with the set of RM-systems $i$ and infected with a plasmids are denoted by $b_i^I$.

**Bacterial population size** The bacterial growth is modeled as in the basic RM-model (section 1.1). In the context of TA-systems, our focus is exclusively on costly plasmids, excluding consideration of plasmids enabling the usage of new energy resources. The density-dependent growth rate for plasmid-free and plasmid-carrying cells is, therefore, given by $\rho_i \left(1 - \frac{B+B^I}{K}\right)$ and $\rho_i\,(1-c)\left(1 - \frac{B+B^I}{K}\right)$, respectively. $\rho_i$ is the maximal division rate of bacterial species $i$, $K$ the carrying capacity, and $B = \sum_{i=1}^{n} b_i$ and $B^I = \sum_{i=1}^{n} b_i^I$ the total bacterial cell density of plasmid free and plasmid carrying cells respectively.

**Segregation loss and conjugation** As we assume that the plasmid carries a TA-system with a lethal effect, if the plasmid is lost at cell division with a probability of $\tau$, the bacterial cell dies unless the TA-system fails. TA failure occurs with a probability of $\epsilon$. In the case where the donor and recipient carry the same set of RM-Systems $i$, conjugation occurs at a rate of $\gamma$ as the plasmid already has the corresponding methylation pattern. Conjugation between bacterial cells of different types of RM-systems occurs rarely at a rate of $\omega_i\gamma$. Additionally, we assume that conjugation to plasmid-free cells can lead to cell death, but only if the TA-system does not fail and if the TA-system is expressed before RM cleavage. The probability that the TA-system is expressed before RM cleavage is denoted by $\theta$. The death rate for plasmid-free cells through conjugation is therefore given by $\gamma\theta\,(1-\epsilon)\,(1-\omega_i)$.

The described processes can be depicted in a schematic flow diagram (figure S5) and captured by the following equations:

$$
\begin{aligned}
\frac{\mathrm{d}b^i}{\mathrm{d}t} =& \rho_i b^i \left(1 - \frac{B+B_I}{K}\right) + \tau_i \epsilon \rho_i\,(1-c_i)\,b_i \left(1 - \frac{B_I + \chi B}{K_I}\right) - \left(\sum_{\substack{j=1\\ j\neq i}}^{n} \gamma\omega_i b_I^j + \gamma b_I^i\right) b^i - \theta\,(1-\epsilon) \sum_{\substack{j=1\\ j\neq i}}^{n} \gamma\,(1-\omega_i)\, b_I^j b^i - d_i b_i,\\
\frac{\mathrm{d}b_I^i}{\mathrm{d}t} =& \rho_i\,(1-c_i)\,(1-\tau_i)\,b_I^i \left(1 - \frac{B_I+\chi B}{K_I}\right) + \left(\sum_{\substack{j=1\\ j\neq i}}^{n} \omega_i \gamma b_I^j + \gamma b_I^i\right) b^i - d_i b_i^I .
\end{aligned}
\tag{2}
$$

| | |
|---|---|
| $b_i$ | **Cells with unique set of RM-system $i$** |
| $b_i^I$ | **Cells with unique set of RM-system $i$ and a plasmid with corresponding methylation** |

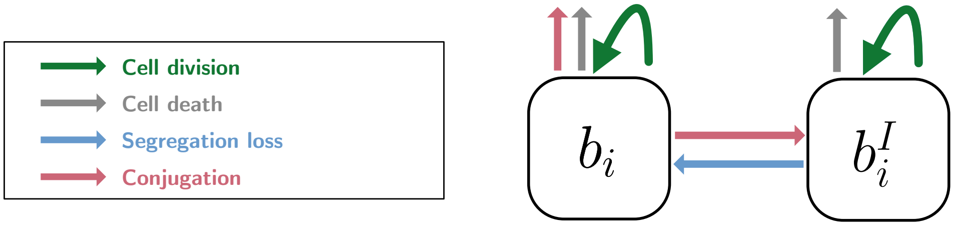


**Figure S5. Visualization of the modeled bacterial population, infection and immunity dynamics of RM and a conjugative plasmid with a lethal TA-system.**

The extension of the RM-model with TA-systems was implemented along the lines of the basic RM-model (section 1.1).

### 2 CRISPR-Cas

#### 2.1 Basic model

**Bacterial population composition** We consider a clonal bacterial population where all cells carry a CRISPR-Cas system. Additionally, we consider a conjugative plasmid that is able to infect bacterial cells that have a CIRSPR-Cas system that is naive to the conjugative plasmid $S$. Once the naive bacterial cells are infected with the plasmid they are denoted by $I$. In case the CRISPR-Cas system has acquired a spacer against the plasmid, and is therefore immune against plasmid infection, the bacterial cell is denoted by $R$. We assume that there is selective pressure to maintain the CRISPR-Cas system by other mobile genetic elements such as phages and therefore do not consider possible loss of CRISPR-Cas systems. Therefore the total bacterial population consists out of three types of cells: $S, I, R$.

**Bacterial population size** The population growth respectively death dynamics are modeled in all our models the same. The bacterial growth is modeled by a density dependent division rate $\rho\left(1-\frac{S+I+R}{K}\right)$, where $\rho$ is the maximal division rate of bacterial species and $K$ the carrying capacity. Plasmids can not only carry beneficial accessory genes that encode resistances against antibiotics or heavy metals, but it has also been reported that plasmids encode genes, which allow the usage of new energy resources such as lactose, other sugar derivatives or light energy [1–3]. We assume that the conjugative plasmid can enable the use of a new energy source, which is reflected in our model with a higher carrying capacity $K_I$ for bacterial cells that carry the plasmid than for cells that do not carry a plasmid ($K_I > K$). Plasmid carrying cells can use the same resources as plasmid free cells and additionally another resource. The factor $\chi$ reflects how much plasmid free cells $S, R$ participate in the resources of plasmid carrying cells $I$. The density dependent growth rate for plasmid carrying cells is therefore described by: $\rho\,(1-c)\left(1-\frac{I+\chi(S+R)}{K_I}\right)$, where $c$ denotes the fitness effect of a plasmid ($c>0$: cost, $c<0$: benefit). We assume that bacterial cells die with a constant rate $d$. Note that CRISPR-Cas immunity can be associated with autoimmunity costs [8], which can be modeled with a lower maximal division rate $\rho$ or a higher death rate $d$. As all types of our cells carry a CRISPR-Cas system this can be directly included into the rates $\rho$ or $d$ such that no extra variable needs to be introduced.

**Segregation loss and conjugation** Cell division of plasmid carrying cells $I$ may not necessary lead to two plasmid carrying daughter cells. At cell division the plasmid gets lost from one daughter cell with probability $\tau$, which is called segregation loss. We assume a well mixed bacterial population, where plasmid conjugation occurs between plasmid free cells $S$ and plasmid carrying $I$ in a density-dependent manner with conjugation rate $\gamma$.

**Spacer acquisition** Given that spacer acquisition has been linked to DNA replication [9], we assume that spacer acquisition from the plasmid can occur whenever the plasmid is replicated, i.e., during both cell division and conjugation. We denote the probability to acquire a spacer at cell division as $\alpha_\rho$ and the probability to acquire a spacer at conjugation with $\alpha_\gamma$.

The described processes can be depicted in a schematic flow diagram (main text figure 1B,C) and captured by the following equations:

$$\begin{aligned}
\frac{\mathrm{d}S}{\mathrm{d}t} &= \rho S\left(1-\frac{S+I+R}{K}\right)+\tau\rho\,(1-c)\,(1-\alpha_\rho)\,I\left(1-\frac{I+\chi\,(S+R)}{K_I}\right)-\gamma SI-dS,\\
\frac{\mathrm{d}I}{\mathrm{d}t} &= \rho\,(1-c)\,(1-\alpha_\rho)\,(1-\tau)\,I\left(1-\frac{I+\chi\,(S+R)}{K_I}\right)+(1-\alpha_\gamma)\,\gamma SI-dI,\\
\frac{\mathrm{d}R}{\mathrm{d}t} &= \rho R\left(1-\frac{S+I+R}{K}\right)+\rho\,(1-c)\,\alpha_\rho I\left(1-\frac{S+I+R}{K}\right)+\alpha_\gamma\gamma SI-dR.
\end{aligned} \tag{3}$$

##### 2.1.1 Important model assumption about CRISPR-Cas

For phages, the probability to acquire a spacer has been estimated to be in the magnitude of $10^{-6}$ [10] . To our knowledge, there is currently no estimate for plasmids. Our basic CRISPR-Cas model is a simple model, not considering the co-evolution between CRISPR-Cas and plasmids. Plasmids can evolve and potentially escape a CRISPR-Cas spacer and therewith immunity. In the case of phages it is known that spacer heterogeneity can be essential to combat phage evolution and drive the phage eventually to extinction [11, 12]. We explore co-evolutionary dynamics between plasmids and CRISPR-Cas in section 2.3.

##### 2.1.2 Implementation

We deterministically simulate the set of differential equations 3 with JULIA. Throughout all simulations, we make the following assumptions: the maximal division rate is set to $\rho = 1.3$, and the death rate to $d = 0.1$. The carrying capacity is set to $K = 10^{10}$. Additionally, unless explicitly stated otherwise, we assume that the plasmid does not confer the ability to utilize an additional resource, meaning $K = K_I$ and $\chi = 1$. We initialize our simulations at maximal population size with plasmid-free cells naive to the plasmid $S$, except for a small proportion of plasmid-infected cells $I = 10^4 \frac{\mathrm{CFU}}{\mathrm{ml}}$. The simulations are run for a maximum of $10^6$ hours. If equilibrium is reached before this time, the simulations are terminated early using the built-in *TerminateSteadyState* callback in the *DifferentialEquations* package [13]. Apart from the global sensitivity analysis discussed in section 2.1.7, we assume that the probability to acquire a spacer at cell division and the probability to acquire a spacer at conjugation are identical (i.e., $\alpha_\rho = \alpha_\gamma$).

##### 2.1.3 Analytical solution of a specific scenario of the basic CRISPR-Cas model

To perform a linear stability analysis of the basic CRISPR-Cas model, as described in the set of differential equations 3, is challenging because it results in obtaining eigenvalues that equal zero. However, we get insights into the dynamics of the basic CRISPR-Cas from linear stability analysis if we simplify the model. In the simplified model we assume that the bacterial population consists of two types of cells: bacterial cells $I$ with a CRISPR-Cas system naive to the plasmid, but infected with a plasmid and bacterial cells that acquired a spacer against the plasmid and are therefore resistant against plasmid infection. Importantly, we assume the plasmid is stably inherited by daughter cells without any segregation loss. The basic CRISPR-Cas model described in the set of differential equations 3 therefore reduces as follows:

$$\begin{aligned}\frac{\mathrm{d}I}{\mathrm{d}t} &= \rho\left(1-c\right)\left(1-\alpha_\rho\right) I \left(1 - \frac{I+\chi R}{K_I}\right) - dI,\\ \frac{\mathrm{d}R}{\mathrm{d}t} &= \rho R\left(1-\frac{I+R}{K}\right) + \rho\left(1-c\right)\alpha_\rho I\left(1-\frac{I+R}{K}\right) - dR.\end{aligned} \tag{4}$$

We assume the plasmid does not enable the bacterial cell to utilize additional resources, therefore $K = K_I$ and $\chi = 1$. First, we apply the negative criterion of Bendixson-Dulac [14] to ensure that the system does not have any periodic orbits and tends to a stationary point. To apply the criterion of Bendixson-Dulac we calculate the divergence of the vector field scaled by $\frac{1}{IR}$ (as $I, R > 0$) and obtain:

$$\begin{aligned}&\frac{\partial}{\partial I}\left[\frac{1}{IR}\left(\rho\left(1-c\right)\left(1-\alpha_\rho\right) I\left(1-\frac{I+R}{K}\right) - dI\right)\right]\\ &+\frac{\partial}{\partial R}\left[\frac{1}{IR}\left(\rho R\left(1-\frac{I+R}{K}\right) + \rho\left(1-c\right)\alpha_\rho I\left(1-\frac{I+R}{K}\right) - dR\right)\right]\\ &= -\frac{b\left(\left(1-c\right)I+R\right)}{KRI} < 0.\end{aligned}$$

As the result is negative, we can exclude according to the criterion of Bendixson-Dulac any periodic orbits and can conclude that the solution tends to a stationary point. Next, we calculate the stationary points. For this, we set the set of differential equations to zero with $K = K_I$ and $\chi = 1$ and obtain three equilibria:

$$E_1 = (I_1, R_1) = (0,0)\,,$$
$$E_2 = (I_2, R_2) = \left(0, K - \frac{dK}{\rho}\right),$$
$$E_3 = (I_3, R_3) = \left(K\left(\frac{d\left(1-\frac{1}{(c-1)(\alpha_\rho-1)}\right)}{bc} - \left(1-\frac{1}{c}\right)\alpha_\rho\right), \frac{K\alpha_\rho\left(\rho-\rho c - d - \rho\left(1-c\right)\alpha_\rho\right)}{\rho c\left(\alpha_\rho-1\right)}\right).$$

To assess the stability of equilibria, we compute the Jacobian matrix of the simplified model described in set of equations 4 at each equilibrium point and determine the corresponding eigenvalues. An equilibrium is linearly stable if all the eigenvalues have a negative real part [14].

The eigenvalues for $E_1$ are given by $\lambda_1^1 = \rho - d$ and $\lambda_2^1 = \rho\left(1-c\right) - d - \rho\alpha_\rho + \alpha_\rho\rho c$. As we assume that the maximal growth rate is bigger than the death rate, i.e. $\rho > d$, the first eigenvalue $\lambda_1^1$ is positive and therefore the first equilibrium $E_1$ is not stable.

The eigenvalues for the second equilibrium $E_2$ are given by $\lambda_1^2 = -\rho - d$ and $\lambda_2^2 = d\left(\rho\alpha_\rho c - c - \alpha_\rho\right)$.

The eigenvalues for the third equilibrium $E_3$ are given by $\lambda_1^3 = \frac{\left(c^2-c\right)\left(\rho-\rho c-d-b\alpha_1+\rho c\alpha_\rho\right)}{c(1-c)}$ and $\lambda_3^2 = \frac{c^2\rho+c\rho\alpha_\rho-c^2 d\alpha_\rho}{c(1-c)(1-\alpha_\rho)}$.

We obtain that $E_2$ has only negative eigenvalues and is therefore stable when $\frac{-\alpha_\rho}{1-\alpha_\rho} < c$ and $E_3$ when $c < \frac{-\alpha_\rho}{1-\alpha_\rho}$. In conclusion, a plasmid-infected population can only persist if the plasmid fitness effect $c$ is negative, representing a benefit in terms of growth, and if this fitness effect can compensate for the spacer acquisition obtained at cell replication.

##### 2.1.4 The maintenance of beneficial plasmids is robust to variations in plasmid traits and strength of CRISPR-Cas immunity

To investigate whether beneficial plasmids are robustly maintained in the presence of CRISPR-Cas, we ran 1000 simulations, varying the extent of plasmid benefit on bacterial cell division $c$, the probability of losing a plasmid $\tau$, the conjugation rate $\gamma$, and the probability of spacer acquisition at cell division $\alpha_\rho$ and at conjugation $\alpha_\gamma$. For parameter variation, we employed Latin hypercube sampling, sampling all parameters on a logarithmic scale, except for the plasmid's fitness effect. The parameter bounds were set as follows: $c = [-2, 0]$, $\tau = [-5, -1]$, $\gamma = [-20, -7]$, $\alpha_\rho = [-20, -4]$, $\alpha_\gamma = [-20, -4]$. We find robust maintenance of beneficial plasmids in 996 out of 1000 simulations (see Figure S6). The four simulations with no plasmid survival share common characteristics: notably low plasmid benefit ($c > -0.05$), high segregation loss, high spacer acquisition probabilities and rather low conjugation rates (Table S6). Experimental evidence suggests that plasmids typically have segregation loss probabilities of at most $10^{-3}$ or lower [15,16]. However, in all four parameter sets, segregation loss is higher than $\tau > 10^{-2}$. Similarly, when the conjugation machinery is fully expressed, plasmid conjugation rates often are higher than $\gamma > 10^{-12}$ [17], but in these four parameter sets, the conjugation rate is below $\gamma < 10^{-14}$. Moreover, known estimates of spacer acquisition rates, derived from phages, are generally low, around $10^{-6}$ [10], yet at least one spacer acquisition rate in these parameter-set is higher than $10^{-5}$. In those four parameter sets, the low plasmid benefit and rather low conjugation rate are insufficient to counterbalance the effects of high segregation loss and high spacer acquisition.

##### 2.1.5 Beneficial plasmids can invade a bacterial population with a high proportion of resistant cells

When investigating the plasmid's ability to invade and survive in a bacterial population, in dependence on the plasmid's fitness effect on cell replication and the initial proportion of resistant cells, we

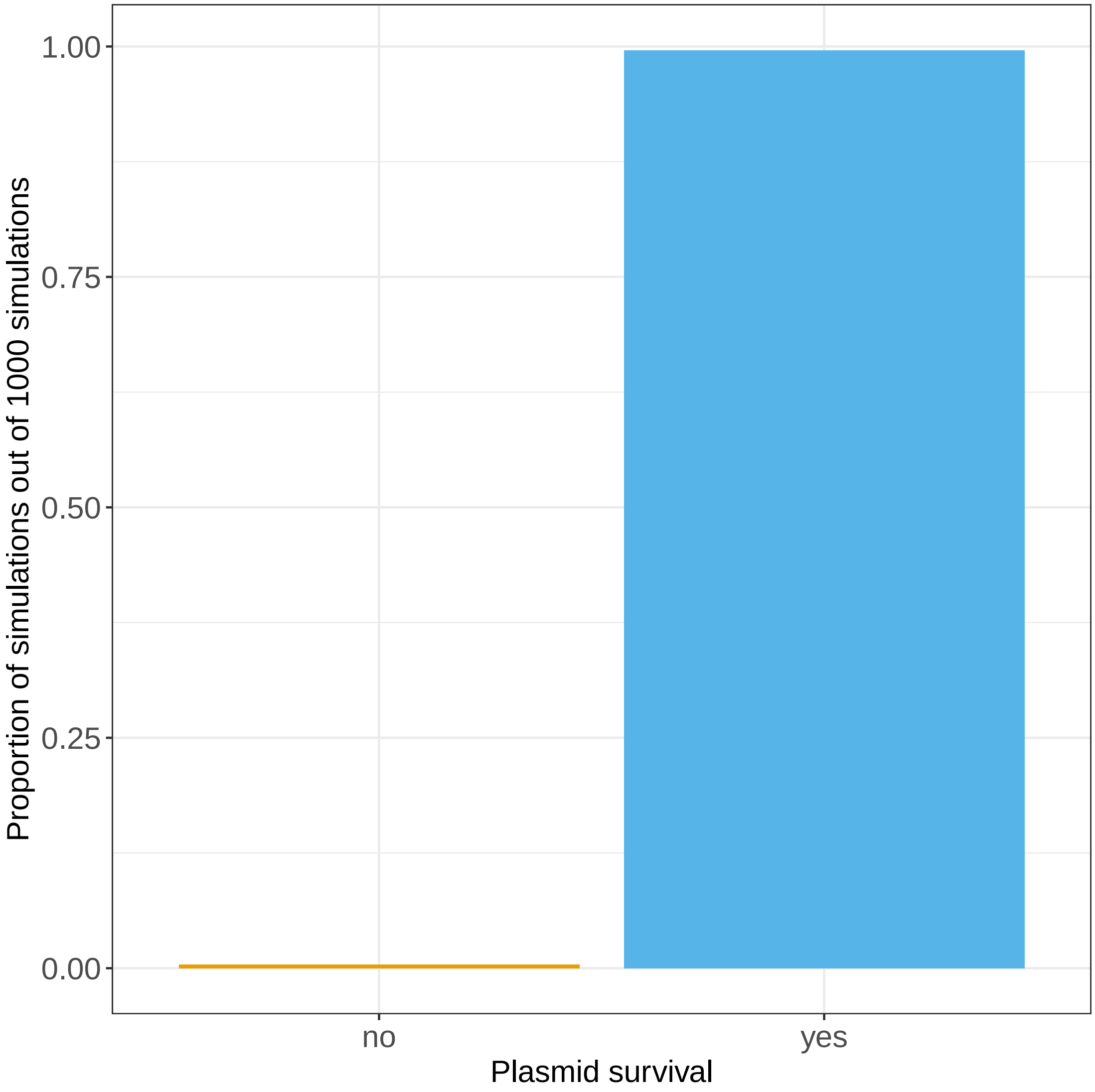


**Figure S6. Beneficial plasmids are robustly maintained in the presence of CRISPR-Cas.** Among 1000 simulations, where the degree of the plasmid's benefit on cell division ($c$), the probability of losing the plasmids ($\tau$), the conjugation rate ($\gamma$), and the probabilities of acquiring a spacer at cell division ($\alpha_\rho$) and at conjugation ($\alpha_\gamma$) were varied using Latin hypercube sampling, 996 simulations demonstrate the persistent maintenance of beneficial plasmids.

observe that beneficial plasmids can be maintained in the bacterial population, regardless of the initial proportion of resistant cells. However, costly plasmids will be excluded in the long term (figure S7).

###### 2.1.6 Plasmids allowing the utilization of an additional resource

When investigating the plasmid's ability to invade and survive in a bacterial population, in dependence on the plasmid's fitness effect on cell replication and the increase in carrying capacity through the plasmid, we find that plasmids enabling the utilization of an additional resource can promote plasmid survival, even when the maximal cell division is reduced due to the plasmid (figure S8). Despite the plasmid's impact on reducing maximal cell division, utilizing the additional resource provides an advantage, especially at high densities when the commonly used resource faces high competition among both plasmid-free and plasmid-carrying cells, leading to increased net-cell growth in plasmid-carrying

| simulation number | $c$ | $\tau$ | $\gamma$ | $\alpha_\rho$ | $\alpha_\gamma$ |
|---|---|---|---|---|---|
| 1 | $-0.024$ | $10^{-1.124}$ | $10^{14.735}$ | $10^{-4.462}$ | $10^{-4.528}$ |
| 2 | $-0.046$ | $10^{-1.316}$ | $10^{-14.254}$ | $10^{-7.528}$ | $10^{-5.446}$ |
| 3 | $-0.004$ | $10^{-1.688}$ | $10^{-14.488}$ | $10^{-4.78}$ | $10^{-5.698}$ |
| 4 | 0 | $10^{-2.488}$ | $10^{-14.215}$ | $10^{-8.2}$ | $10^{-4.072}$ |

**Table S1.** Overview of the four parameter-sets where beneficial plasmids do not lead to the maintenance of the plasmid. $c$ denotes the plasmid's fitness effect on cell replication, $\tau$ the probability to lose the plasmid, $\gamma$ the conjugation rate and $\alpha_\rho$ and $\alpha_\gamma$) the probabilities to acquire a spacer at cell replication and conjugation, respectively. Note that in simulation 4, the plasmid does not confer any benefit or cost, but its effect on cell replication is rather neutral.

cells.

###### 2.1.7 Global sensitivity analysis assessing which parameters determine the speed with which costly plasmids go to extinction

To assess which parameters are most important to determine how fast a costly plasmid is driven to extinction we performed a global sensitivity analysis with the variance based Sobol's method. For the implementation of the Sobol's method we used the *GlobalSensitivity* (version 2.1.4) [18] and used the total order Sobol's indices to assess the importance of parameters. Unlike the first-order Sobol's index, which measures the effect of varying a single parameter in isolation, the total order Sobol's index considers not only the individual parameter's impact but also its contribution to output variance through interactions with other parameters. It is important to note that the sum of Sobol's indices may be larger than one, as interactions of parameters are accounted for in several individual Sobol's indices.

The calculation of total order Sobol's indices was based on 1000 simulations with varying parameter combinations. We varied the following parameters and sampled them from the specified ranges: the plasmid effect on bacerial cell division $c \in [0.05, 0.2]$, probability of losing the plasmid $\tau \in [10^{-5}, 10^{-3}]$, conjugation rate $\gamma \in [10^{-11}, 10^{-7}]$, probability of spacer acquisition at cell division $\alpha_\rho \in [10^{-10}, 10^{-4}]$ and probability of spacer acquisition at conjugation $\alpha_\gamma \in [10^{-10}, 10^{-4}]$. Note that we restricted our ranges to investigate cases where the costly plasmid is able to maintain itself without CRISPR-Cas.

##### 2.2 TA-systems

**Bacterial population composition** The bacterial population consists of the same bacterial cells as in the basic CRISPR-Cas and plasmid model (see section 2.1). Bacterial cells with a naive CRISPR-Cas system to the plasmid without a plasmid are denoted by $S$. Cells carrying a plasmid and a naive CRISPR system are denoted by $I$. Bacterial cells that acquired a spacer against the plasmid, and are therewith immune, are denoted by $R$. The plasmid carries a TA-system, which we assume has an lethal effect for the bacterial cell without a the anti-toxin.

**Bacterial population size** The population growth is modeled as in the basic CRISPR-Cas model (see section 2.1). In the context of TA-systems, our focus is exclusively on costly plasmids, excluding consideration of plasmids enabling the usage of new energy resources. The density-dependent growth rate for plasmid-free and plasmid-carrying cells is, therefore, given by $\rho S\left(1-\frac{T}{K}\right)$ and $\rho S\left(1-c\right)\left(1-\frac{T}{K}\right)$, respectively. Here, $T := S + I + R$ denotes the total population size, $\rho$ represents the maximal cell division rate, $K$ is the carrying capacity, and $c$ signifies the plasmid's fitness effect on cell replication. We assume that cells die at a constant rate $d$.

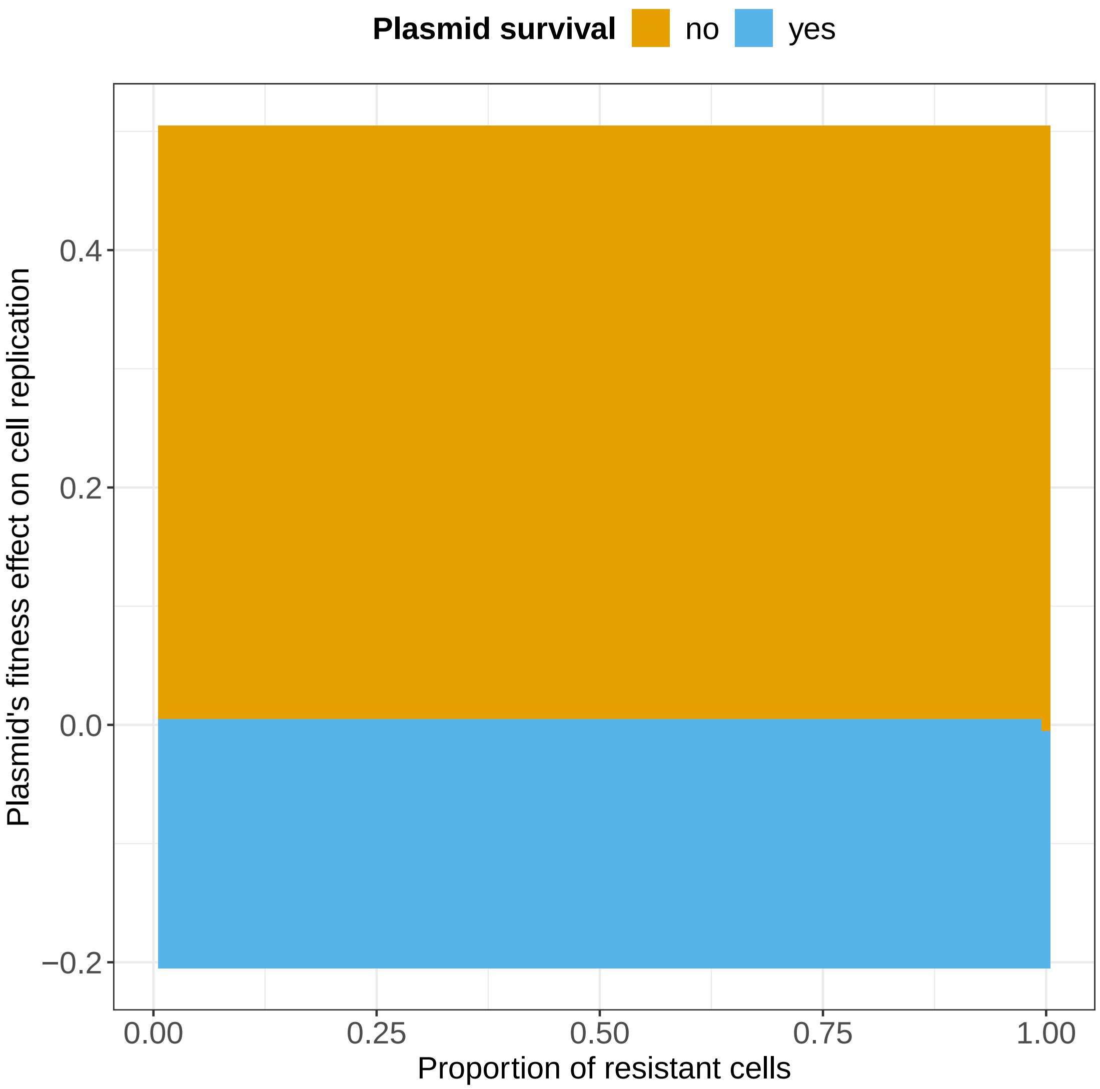


**Figure S7. Plasmid survival: dependency on the plasmid's fitness effect on cell replication and proportion of initial resistant cells.** The ability of a plasmid to survive upon entering a bacterial population, depending on the plasmid's fitness effect on cell replication and the initial proportion of resistant cells in the population, is shown. Negative values of $c$, representing the plasmid's fitness effect on cell replication, indicate beneficial plasmids, whereas positive values indicate costly plasmids. Only beneficial plasmids are maintained, independent of the initial proportion of resistant cells. In the simulations, we set the conjugation rate to $\gamma = 10^{-10}$ and the probability of losing a plasmid to $\tau = 10^{-4}$.

**Segregation loss and conjugation** As we assume that the plasmid carries a TA-system with a lethal effect, if the plasmid is lost at cell division with probability $\tau$, the bacterial cell dies unless the TA-system fails. We assume that the TA-system fails with probability $\epsilon$. Conjugation is modeled in a density dependent manner with conjugation rate $\gamma$. We assume that when a plasmid enters a resistant cell $R$ through conjugation, the plasmid's TA-system might be expressed before CRISPR-Cas interference, with a probability denoted as $\theta_\gamma$. Consequently, if a resistant cell receives a plasmid, it might result in cell death if the TA-system does not fail. The death rate for resistant cells $R$ through conjugation is given by $\theta_\gamma(1-\epsilon)\gamma$.

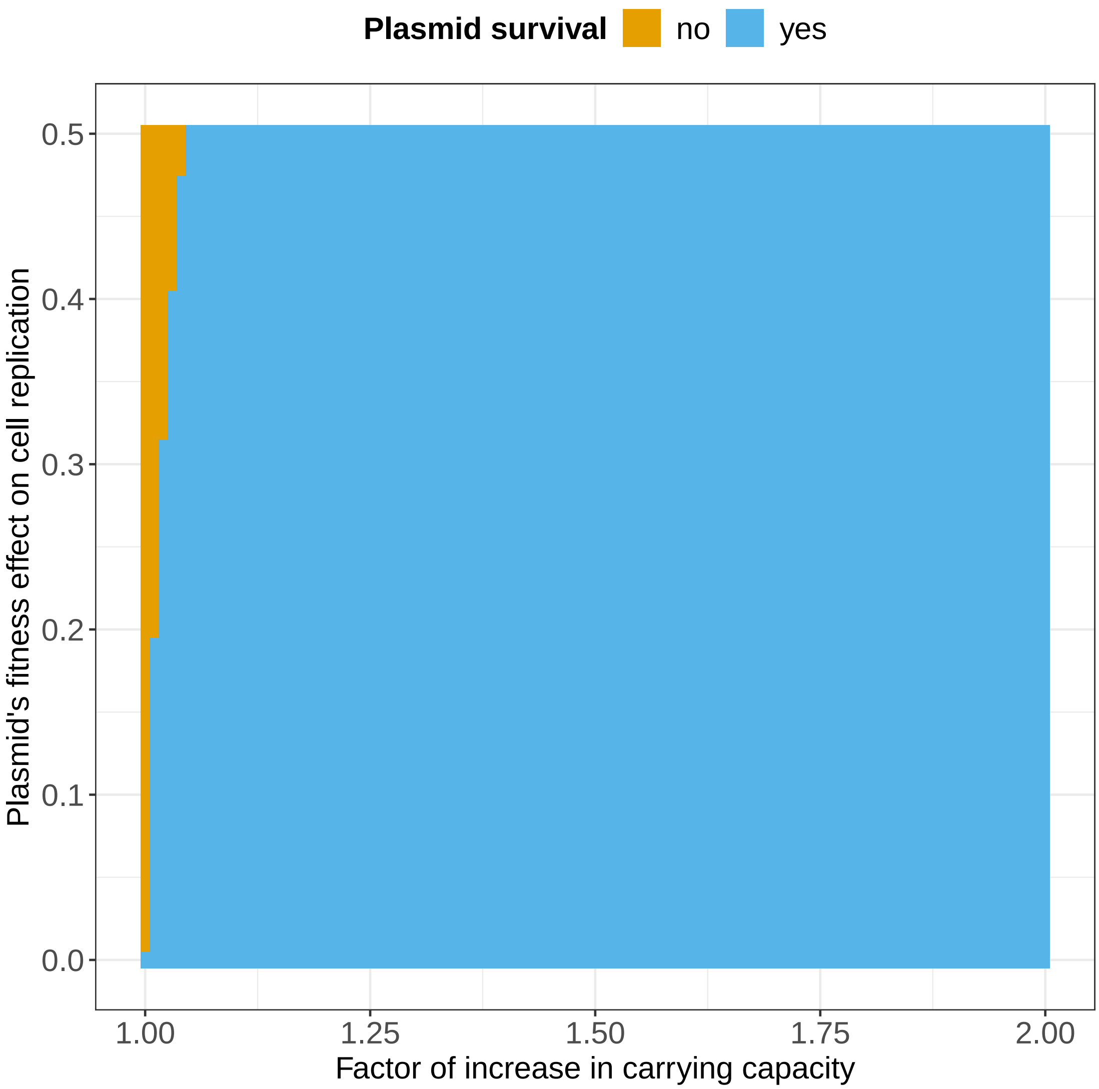


**Figure S8. Plasmid survival: dependency on the plasmid's fitness effect on cell replication and the increase in carrying capacity through the plasmid.** The ability of a plasmid to survive upon entering a bacterial population, depending on the plasmid's fitness effect on cell replication and the increase in carrying capacity through the plasmid, is shown. The factor of increase, denoted as $\iota$, represents the multiplication factor by which the carrying capacity $K_I$ exceeds that of non-plasmid-infected cells ($K_I = \iota K$). $\chi$ the factor indicating the extent of participation of plasmid-free cells in the resources of plasmid-carrying cells is given by $\chi = \frac{K}{K_I}$. In the simulations, we set the conjugation rate to $\gamma = 10^{-10}$ and the probability of losing a plasmid to $\tau = 10^{-4}$.

**Spacer acquisition** We assume spacer acquisition of the plasmid can occur during plasmid replication, specifically at cell division and conjugation. The probability of acquiring a spacer at cell division is denoted as $\alpha_\rho$, while the probability at conjugation is denoted as $\alpha_\gamma$. However, if the TA-system is expressed before CRISPR-Cas interference and does not fail, it can lead to the death of cells that newly acquired a spacer. We denote the probability that the TA-system was expressed before CRISPR-Cas interference for cells that newly acquired a spacer as $\theta_\alpha$. We assume that the probabilities of TA-system failure $\epsilon$ and TA-system expression before CRISPR-Cas interference $\theta_\alpha$ are independent.

The described processes can be depicted in a schematic flow diagram (figure S9) and captured by the following equations:

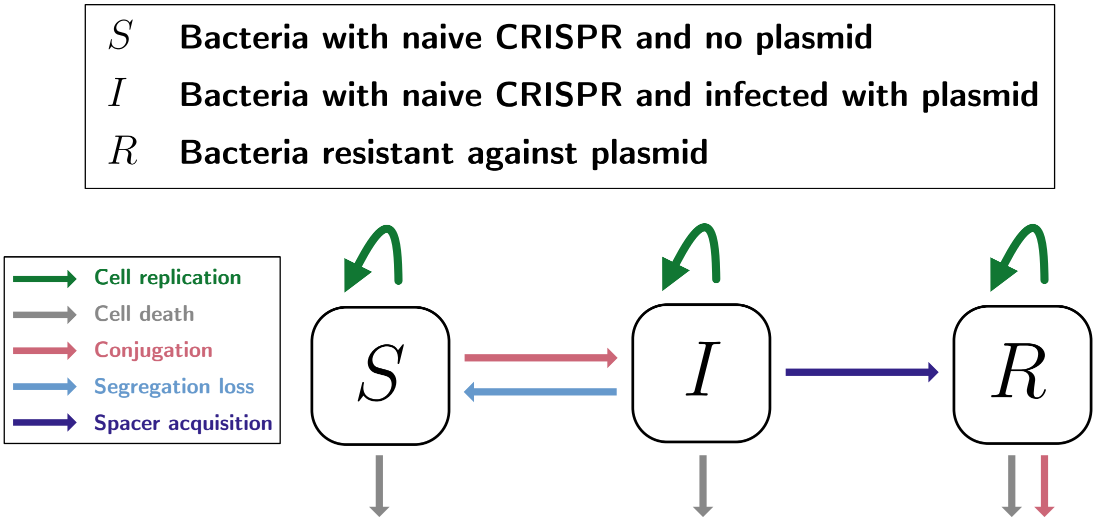


**Figure S9. Visualization of the modeled bacterial population, infection and immunity dynamics of CRISPR-Cas and a conjugative plasmid with a lethal TA-system.**

$$
\begin{aligned}
\frac{\mathrm{d}S}{\mathrm{d}t} &= \rho S\left(1-\frac{T}{K}\right) + \tau\epsilon\rho\,(1-c)\,(1-\alpha_b)\,I\left(1-\frac{T}{K}\right) - \gamma SI - dS \\
\frac{\mathrm{d}I}{\mathrm{d}t} &= \rho\,(1-c)\,(1-\alpha_b)\,(1-\tau)\,I\left(1-\frac{T}{K}\right) + (1-\alpha_\gamma)\,\gamma SI + -dI \\
\frac{\mathrm{d}R}{\mathrm{d}t} &= \rho R\left(1-\frac{T}{K}\right) + (\epsilon + (1-\theta_\alpha) - \epsilon\theta_\alpha)\,\rho\,(1-c)\,\alpha_b I\left(1-\frac{T}{K}\right) + (\epsilon + (1-\theta_\alpha) - \epsilon\theta_\alpha)\,\alpha_\gamma\gamma SI - \theta_\gamma\,(1-\epsilon)\,\gamma RI - dR
\end{aligned}
\tag{5}
$$

The extension of the basic CRISPR-Cas model with TA-systems was implemented along the lines of the basic CRISPR-Cas model (section 2.1).

#### 2.3 CRISPR-Cas and plasmid co-evolution

In our main CRISPR-Cas model we find that costly plasmids are eliminated from the population by CRISPR-Cas. However, it is important to note that in our basic model we make the assumption that the plasmid can't escape a spacer by mutating its protospacer, while it is well known that (i) phages can frequently escape spacers by mutation [19] and (ii) that plasmids can mutate and undergo evolutionary change [20]. Yet, when a virulent phage faces CRISPR-Cas immunity, the successful escape from a single spacer does not guarantee the phage's survival. The number of potential protospacers in the phage genome can vary, ranging up to several hundreds [21, 22] and the choice among them is stochastic (with some biases) [23, 24]. When a virulent phage enters a bacterial population defending using CRISPR-Cas, each cell can acquire independently a spacer, which results in a population-wide spacer diversity. This population-wide spacer diversity, if high enough, can lead the phage to extinction [11, 12, 22]. Alternatively, if the phage is potent enough or the population-wide spacer diversity is too low, the phage can (i) either drive the bacterial population to extinction [12] or (ii) result in a co-evolution, where bacteria acquire multiple spacers [23, 25].

To assess whether CRISPR-Cas can drive costly plasmids to extinction when plasmids can escape spacers trough point mutation, we formulated a mathematical model taking into account some of the co-evolution between CRISPR-Cas and plasmids. To take a first step into investigating the evolutionary dynamics between CRISPR-Cas and plasmids, we consider that each bacterial cell can acquire a single spacer from the plasmid, thereby incorporating population-wide spacer heterogeneity. Additionally, we assume that plasmids can escape a spacer through mutations in their protospacer. However, our model does not account for the possibility that cells can acquire spacers from a mutated protospacer, nor does it consider that a single bacterial cell can acquire multiple distinct plasmid spacers. As a result, we neglect within-cell spacer heterogeneity, which is also known to be important for long-term evolution between phages and CRISPR-Cas.

**Bacterial population composition** We consider a clonal bacterial population where all cells carry a CRISPR-Cas system. Additionally, we consider a conjugative plasmid able to infect bacterial cells if they do not have a corresponding spacer yet. The wild type ($WT$) plasmid, without any modifications due to mutation, can infect bacterial cells with a naive CRISPR-Cas system $S$. Once a bacterial cell is infected with the $WT$ plasmid, it is denoted as $I_{WT}$. We assume that the plasmid has $n_s$ protospacers. In case one of the protospacers $i \in \{1 \dots n_s\}$ mutates, the cell infected with the mutated plasmid with mutation in protospacer $i$ is denoted by $R^i$. A resistant cell with spacer $i$ can still be infected by plasmids that have a mutation in the corresponding protospacer; these cells are denoted by $I_i R^i$.

**Bacterial population size** The population growth is modeled as in the basic CRISPR-Cas model (see section 2.1). In the context of co-evolution between CRISPR-Cas and plasmids, our focus is exclusively on costly plasmids, excluding consideration of plasmids enabling the usage of new energy resources. The density-dependent growth rate for plasmid-free and plasmid-carrying cells is, therefore, given by $\rho S \left(1 - \frac{T}{K}\right)$ and $\rho S \left(1 - c_{wt}\right)\left(1 - \frac{T}{K}\right)$, respectively. Here, $T \coloneqq S + I_{WT} + \sum_{i=1}^{n_s} I_i + \sum_{i=1}^{n_s} R_i + \sum_{i=1}^{n_s} I_i R^i$ denotes the total population size, $\rho$ represents the maximal cell division rate, $K$ is the carrying capacity, and $c_{wt}$ signifies the $WT$ plasmid's fitness effect on cell replication. We assume that the mutation of the $WT$ plasmid results in an altered fitness effect, such that the escape plasmid has the plasmid fitness effect $c_e$. Therefore, the growth rate for mutated plasmids is given by $\rho S \left(1 - c_e\right)\left(1 - \frac{T}{K}\right)$. We also assume that cells die at a constant rate $d$.

**Segregation loss and conjugation** Cell division of plasmid-carrying cells, such as $I_{WT}$, $I_i$, or $I_i R^i$, may not necessarily lead to two plasmid-carrying daughter cells. During cell division, the plasmid has a probability of $\tau$ to be lost from one of the daughter cells. We assume a well-mixed bacterial population, wherein plasmid conjugation occurs between plasmid-free cells and plasmid-carrying cells in a density-dependent manner. The conjugation rate is denoted as $\gamma_{wt}$ for $WT$ plasmids and $\gamma_e$ for escape plasmids with mutations. The $WT$ plasmid can only conjugate into naive cells $S$, while the mutated plasmid with a mutation in protospacer $i$ can conjugate into both naive cells $S$ and cells that carry spacer $i$, from which the mutated plasmid escapes.

**Spacer acquisition** We assume that spacer acquisition from plasmids can occur whenever the plasmid is replicated, i.e. during both cell division and conjugation. We denote the probability to acquire any spacer at cell division as $\alpha_\rho$ and the probability to acquire any spacer at conjugation with $\alpha_\gamma$.

**Plasmid spacer escape** We assume that plasmids have the possibility to mutate whenever they replicate, i.e., during cell replication and conjugation. Specifically, each of the $n_s$ protospacers has a mutation probability of $\mu_\rho$ during cell replication and $\mu_\gamma$ during conjugation. In case a plasmid has a mutation in protospacer $i$ it can escape the corresponding spacer, i.e. the plasmid can infect cells $R^i$.

The described processes can be depicted in a schematic flow diagram (figure S10) and captured by the following equations:

$$
\begin{aligned}
\frac{dS}{dt} ={}& \underbrace{\rho S\left(1-\frac{T}{K}\right)}_{\text{replication}} - \underbrace{dS}_{\text{death}} - \underbrace{(1-\alpha_\gamma)\,(1-n_s\mu_\gamma)\gamma_{wt}SI_{wt}}_{\substack{\text{conjugation without spacer}\\ \text{acquisition and mutation}}} - \underbrace{(1-\alpha_\gamma)n_s\mu_\gamma\gamma_{wt}SI_{wt}}_{\substack{\text{conjugation without spacer}\\ \text{acquisition and with mutation}}} \\
& - \underbrace{(1-\alpha_\gamma)\gamma_e S\left(\sum_{i=1}^{n_s} I_i + \sum_{i=1}^{n_s} I_i R^i\right)}_{\substack{\text{conjugation without spacer}\\ \text{acquisition}}} - \underbrace{\alpha_\gamma\gamma_{wt}SI_{wt}}_{\substack{\text{conjugation with spacer}\\ \text{acquisition}}} \underbrace{- \alpha_\gamma\gamma_e S\left(\sum_{i=1}^{n_s} I_i + \sum_{i=1}^{n_s} I_i R^i\right)}_{\substack{\text{conjugation with spacer}\\ \text{acquisition}}} \\
& + \underbrace{\tau\rho(1-c_{wt})(1-\alpha_\rho)I_{wt}\left(1-\frac{T}{K}\right)}_{\text{segregation loss}} + \underbrace{\tau b(1-c_e)(1-\alpha_\rho)\sum_{i=1}^{n_s} I_i\left(1-\frac{T}{K}\right)}_{\text{segregation loss}}
\end{aligned} \tag{6}
$$

$$
\begin{aligned}
\frac{dI_{wt}}{dt} ={}& \underbrace{\rho(1-c_{wt})I_{wt}\left(1-\frac{T}{K}\right)}_{\text{replication}} - \underbrace{dI_{wt}}_{\text{death}} + \underbrace{(1-\alpha_\gamma)\,(1-n_s\mu_\gamma)\gamma_{wt}SI_{wt}}_{\substack{\text{conjugation without spacer}\\ \text{acquisition and mutation}}} \underbrace{-\alpha_\rho\rho(1-c_{wt})I_{wt}\left(1-\frac{T}{K}\right)}_{\substack{\text{spacer acquisition at}\\ \text{replication}}} \\
& - \underbrace{\tau\rho(1-c_{wt})(1-\alpha_\rho)I_{wt}\left(1-\frac{T}{K}\right)}_{\text{segregation loss}} - \underbrace{n_s\mu_\rho\rho(1-c_{wt})(1-\alpha_\rho)I_{wt}\left(1-\frac{T}{K}\right)}_{\text{protospacer mutation}}
\end{aligned} \tag{7}
$$

$$
\begin{aligned}
\frac{dI_i}{dt} ={}& \underbrace{\rho(1-c_e)I_i\left(1-\frac{T}{K}\right)}_{\text{replication}} - \underbrace{dI_i}_{\text{death}} + \underbrace{(1-\alpha_\gamma)\mu_\gamma\gamma_{wt}SI_{wt}}_{\substack{\text{conjugation without spacer}\\ \text{acquisition and with mutation}}} + \underbrace{(1-\alpha_\gamma)\gamma_e S(I_i + I_iR^i)}_{\substack{\text{conjugation without spacer}\\ \text{acquisition}}} \\
& - \underbrace{\alpha_\rho\rho(1-c_e)I_i\left(1-\frac{T}{K}\right)}_{\substack{\text{spacer acquisition at}\\ \text{replication}}} - \underbrace{\tau\rho(1-c_e)(1-\alpha_b)I_i\left(1-\frac{T}{K}\right)}_{\text{segregation loss}} + \underbrace{\mu_\rho b(1-c_{wt})(1-\alpha_\rho)I_{wt}\left(1-\frac{T}{K}\right)}_{\text{protospacer mutation at replication}}
\end{aligned} \tag{8}
$$

$$
\begin{aligned}
\frac{dR^i}{dt} ={}& \underbrace{\rho R^i\left(1-\frac{T}{K}\right)}_{\text{replication}} - \underbrace{dR^i}_{\text{death}} + \underbrace{\frac{\alpha_\gamma}{n_s}\gamma_{wt}SI_{wt}}_{\substack{\text{conjugation with}\\ \text{acquisition}}} \underbrace{+ \frac{\alpha_\gamma}{n_s-1}\gamma_e S\left(\sum_{\substack{j=1\\ j\neq i}}^{n_s} I_j + \sum_{\substack{j=1\\ j\neq i}}^{n_s} I_jR^j\right)}_{\substack{\text{conjugation with spacer}\\ \text{acquisition}}} \\
& + \underbrace{\frac{\alpha_\rho}{n_s-1}\rho(1-c_e)\sum_{\substack{j=1\\ j\neq i}}^{n_s} I_j\left(1-\frac{T}{K}\right)}_{\substack{\text{spacer acquisition at}\\ \text{replication}}} + \underbrace{\frac{\alpha_\rho}{n_s}\rho(1-c_{wt})I_{wt}\left(1-\frac{T}{K}\right)}_{\substack{\text{spacer acquisition at}\\ \text{replication}}} - \underbrace{\gamma_e R^i(I_i + I_iR^i)}_{\text{conjugation}} \\
& + \underbrace{\tau b(1-c_e)(1-\alpha_\rho)I_iR^i\left(1-\frac{T}{K}\right)}_{\text{segregation loss}}
\end{aligned} \tag{9}
$$

$$
\frac{dI_iR^i}{dt} = \underbrace{\rho I_iR^i\left(1-\frac{T}{K}\right)}_{\text{replication}} - \underbrace{dI_iR^i}_{\text{death}} + \underbrace{\gamma_e R^i(I_i + I_iR^i)}_{\text{conjugation}} \underbrace{- \tau\rho(1-c_e)(1-\alpha_\rho)I_iR^i\left(1-\frac{T}{K}\right)}_{\text{segregation loss}} \tag{10}
$$

**Important model assumptions about the evolution of CRISPR-Cas and plasmids** Generally, estimating plasmid mutation rates is challenging [26]. For protospacer mutation probability at cell replication $\mu_\rho$ and at conjuation $\mu_\gamma$, we choose values of $10^{-8}$ and $10^{-6}$ reflecting a range presented by Deatherage *et al.* [27].

The number of plasmid protospacers $n_s$ can be predicted from the protospacer adjacent motif (PAM) of CRISPR-Cas and the plasmid genome sequence. We predicted the number of protospacers for two plasmids (RP4 [28] and pKJK5 [29]) facing two CRISPR-Cas systems with PAMs of different lengths: the 2-nucleotide PAM of *Escherichia coli* KD263 (AAG) [30] and the 4-nucleotide PAM of

*Streptococcus thermophilus* DGCC7710 (AGAA) [31]. Using the research function of Snapgene Viewer 5.3.1 (from Insightful Science; available at snapgene.com), we predicted the number of protospacers $n_s$ for each of the two PAMs carried by our model plasmids. From the 2-nucleotide PAM, we derived that RP4 and pKJK5 carry, respectively, $n_s = 1823$ and $n_s = 1860$. From the 4-nucleotide PAM, we derived that RP4 carries $n_s = 403$, while pKJK5 carries $n_s = 349$. In our simulations we tested the influence of the number of protospacers with the following values $n_s = \{400, 700, 1000, 1500\}$. We observed that between those values of protospacers the final plasmid prevalence only varies in a small range ($\ll 1\%$) (figure S11). We calculated plasmid prevalence as follows: $\frac{I_{wt}+\sum_{i=1}^{n_s} I_i+\sum_{i=1}^{n_s} I_i R^i}{S+I_{wt}+\sum_{i=1}^{n_s} I_i+\sum_{i=1}^{n_s} I_i R^i+\sum_{i=1}^{n_s} R^i}$. Given the high computational resources needed for simulations with high protospacer numbers and the low variance in output, we used $n_s = 400$ for further simulations.

**Implementation** We stochastically simulate the equations (6– 10) using the *adaptivetau* package in R [32]. The initial conditions for all simulations are set as follows: $S_0 = 8 \times 10^9$, $I_{wt,0} = 10^4$. Throughout all simulations, we maintain a carrying capacity of $K = 10^{10}$, a maximum cell division rate $\rho = 0.55$, a probability to lose the plasmid of 0.05, and a death rate $d = 0.1$. The number of plasmid protoscapers is set to $n_s = 400$ as reasoned in the previous paragraph. Additionally, we assume that the $WT$ plasmid has a plasmid fitness effect on cell replication of $c_{wt} = 0.1$, and escape plasmids with mutations in protospacers have a fitness effect on replication of $c_e = 0.2$. We assume that the conjugation rate is identical for both the $WT$ plasmid and escape plasmids, i.e., $\gamma_{wt} = \gamma_e$. As discussed in the main text, varying assumptions about the effects of mutations on the plasmids, such as the conjugation rate, could result in different model outcomes. All simulations were run for 2000 hours.

#### 2.4 *Bet/Exo*

A newly discovered CRISPR-Cas escape mechanism, *Bet/Exo*, has been identified in IncC plasmids, a clinically relevant family of conjugative plasmids [33]. In contrast to Anti-CRISPR proteins, *Bet/Exo* doesn't prevent interference; instead, it enables plasmids to survive interference by repairing DNA breaks caused during interference and inducing mutations in the protospacer. This process involves utilizing small sequences known as repeats, distributed throughout the plasmid, as primers to initiate RecA-independent homologous recombination [33]. Consequently, *Bet/Exo* induces significant alterations in the protospacer and its surroundings, facilitating escape from CRISPR-Cas. Furthermore, the recombination by *Bet/Exo* is a stochastic process, leading to non-identical repairs of plasmid molecules with the same targeted protospacer [33]. Because (i) CRISPR-Cas spacers often target plasmid conjugation genes [34] and (ii) *Bet/Exo* significantly and stochastically alters the protospacer, the effect on the plasmid's ability to conjugate varies, sometimes resulting in a loss of conjugation ability [33].

##### 2.4.1 Bet-Exo and CRISPR-Cas model

**Bacterial population composition** We consider a bacterial population where all cells carry a CRISPR-Cas system and we consider a conjugative plasmid with a Bet-exo system. Cells that carry a naive CRISPR-Cas system to the wild type plasmid, i.e. no modification of the plasmid through recombination of the *Bet/Exo* system, and are not infected with a plasmid are denoted by $S$. Cells that are naive to the conjugative wild type plasmid and are infected with the wild-type plasmid are denoted by $I$. Cells naive to the conjugative wild-type plasmid but infected with a plasmid that underwent mutation via the *Bet/Exo* system are denoted by $I_m$. In case a bacterial cell has acquired a spacer of the wild-type plasmid and is therewith immune against the wild-type plasmid, it is denoted by $R$. Cells that acquired a spacer against the wild-type plasmid are however not immune against the mutated plasmid and can therefore be infected with a plasmid mutant, those cells are denoted by $R_{I_m}$.

**Bacterial population size** The population growth is modeled following the basic CRISPR-Cas model (see Section 2.1). Since our focus is on costly plasmids in the context of *Bet/Exo*, we omit consideration of plasmids enabling the usage of a new energy resource. The density-dependent cell division rates for plasmid-free cells, plasmid wild-type carrying cells, and plasmid-infected cells with the modified plasmid are therefore given by $\rho\left(1-\frac{T}{K}\right)$, $\rho\left(1-c\right)\left(1-\frac{T}{K}\right)$, and $\rho\left(1-c_m\right)\left(1-\frac{T}{K}\right)$, respectively. Here, $T$ denotes the total bacterial population, given by $T \coloneqq S + I + I_m + R + R_{I_m}$, $K$ the carrying capacity and $\rho$ the maximal division rate. The wildtype plasmid fitness effect on cell division is denoted by $c$ and the mutated ones is denoted by $c_m$.
Any potential cost associated with *Bet/Exo* expression can be factored into the plasmid fitness effects, denoted by $c$ and $c_m$.

**Segregation loss and conjugation** The plasmid dynamics of segregation loss and conjugation are modeled as in the basic CRISPR-Cas and plasmid model (see section 2.1). The conjugation rate of the mutant plasmid is denoted by $\gamma_m$ and the probability of segregation loss is denoted by $\tau_m$.

**Spacer acquisition** Spacer acquisition is modeled identical to the basic CRISPR-Cas and plasmid model (see section 2.1). We assume that only spacers against the wild-type plasmid can be acquired and not the from *Bet/Exo* modified plasmid.

**Plasmid spacer escape** Plasmid carrying *Bet/Exo* can escape plasmid cleavage by repairing double-stranded breaks via recombination [33], the alteration of the plasmid DNA usually lets the plasmid escape the acquired plasmid spacer. We therefore assume that whenever CRISPR-Cas acquires a spacer that only part of the cells can actually degrade the plasmid. With probability $\xi$ we assume that *Bet/Exo* is able to mutate the plasmid accordingly such that CRISPR-Cas immunity is escaped. We assume that the mutation of plasmids can have three distinct effects on plasmid fitness: (i) alteration of the maximal conjugation rate $\gamma$ ($\gamma_m$), (ii) alteration of the probability of segregation loss $\tau$ ($\tau_m$) and (iii) alteration of the plasmid fitness effect on bacterial growth $c$ ($c_m$). Note that in our model we only have one resistant state and do not trace spacer heterogeneity within the bacterial population. We assume that a mutated plasmid can escape CRISPR-Cas immunity. Furthermore, we assume that against the mutated plamid no immunity can be acquired. We therefore do not capture in this simple model the complex dynamics of CRISPR-Cas immunity against escape plasmids and heterogeneity in plasmid mutants and population wide spacer composition.

The described processes can be depicted in a schematic flow diagram (figure S12) and captured by the following equations:

$$
\begin{aligned}
\frac{\mathrm{d}S}{\mathrm{d}t} =& \rho S\left(1-\frac{T}{K}\right) + \left(\tau\rho\left(1-c\right)I + \tau_m\rho\left(1-c_m\right)I_m\right)\left(1-\alpha_\rho\right)\left(1-\frac{T}{K}\right)\\
&- S\left(\gamma I + \gamma_m\left(I_m + R_{I_m}\right)\right) - dS\\
\frac{\mathrm{d}I}{\mathrm{d}t} =& \rho\left(1-c\right)\left(1-\alpha_\rho\right)\left(1-\tau\right)I\left(1-\frac{T}{K}\right) + \left(1-\alpha_\gamma\right)\gamma SI - dI\\
\frac{\mathrm{d}I_m}{\mathrm{d}t} =& \rho\left(1-c_m\right)\left(1-\tau_m\right)I_m\left(1-\frac{T}{K}\right) + \gamma_m S\left(I_m + R_{I_m}\right) - dI\\
\frac{\mathrm{d}R}{\mathrm{d}t} =& \rho R\left(1-\frac{T}{K}\right) + \tau_m\rho\left(1-c_m\right)R\left(1-\frac{T}{K}\right) + \left(1-\xi\right)\rho\left(1-c\right)\alpha_\rho I\left(1-\frac{T}{K}\right)\\
&+ \left(1-\xi\right)\alpha_\gamma\gamma SI - \gamma_m RR_{I_m} - dR\\
\frac{\mathrm{d}R_{I_m}}{\mathrm{d}t} =& \rho\left(1-c_m\right)\left(1-\tau_m\right)R_{I_m}\left(1-\frac{T}{K}\right) + \xi\rho\left(1-c\right)\alpha_\rho I\left(1-\frac{T}{K}\right) + \xi\alpha_\gamma\gamma SI\\
&+ \gamma_m RR_{I_m} - dR_{I_m}
\end{aligned}
\tag{11}
$$

##### 2.4.2 Global sensitivity analysis: identifying key parameters for *Bet/Exo*'s impact on the maintenance of costly plasmids

To assess which parameters are most important in determining whether *Bet/Exo* allows costly plasmids to be maintained, we performed a global sensitivity analysis using the variance-based Sobol's method. For the implementation of the Sobol's method we used the *GlobalSensitivity* (version 2.1.4) [18] and used the total order Sobol's indices to assess the importance of parameters. Unlike the first-order Sobol's index, which measures the effect of varying a single parameter in isolation, the total order Sobol's index considers not only the individual parameter's impact but also its contribution to output variance through interactions with other parameters. It is important to note that the sum of Sobol's indices may be larger than one, as interactions of parameters are accounted for in several individual Sobol's indices.

The calculation of total order Sobol's indices was based on 1000 simulations with varying parameter combinations. We varied the following parameters and sampled them from the specified ranges: the modified plasmid effect on bacterial cell division $c_m \in [0.1, 0.5]$, probability of losing the modified plasmid $\tau_m \in [10^{-4}, 1]$, conjugation rate of the modified plasmid $\gamma_m \in [10^{-20}, 10^{-11}]$, and the probability of *Bet/Exo* modification before CRISPR-Cas interference $\xi \in [0, 1]$. Note that our ranges are constrained to ensure that the modified plasmid is less fit than the wild-type plasmid, under the assumption that negative fitness effects due to modifications by *Bet/Exo* are more probable.

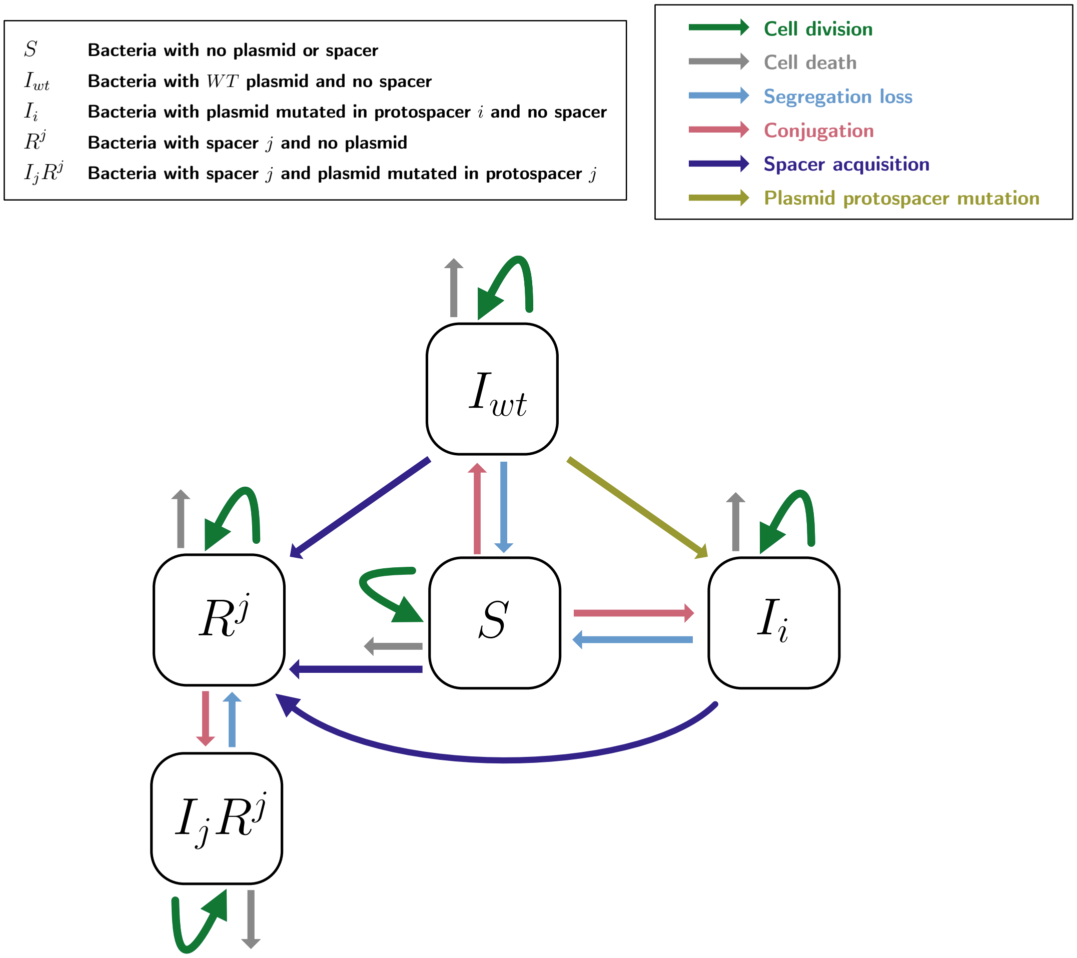


**Figure S10. Visualization of the modeled dynamics encompassing bacterial population, infection, and immunity, considering the co-evolution between CRISPR-Cas and a conjugative plasmid.** Simplified flow diagram of equations (6– 10). In the flow diagram the index $i$ and $j$ are used to illustrate that a naive CRISPR-Cas system infected with an escape plasmid with mutation in protospacer $i$ cannot acquire the mutated spacer $i$, but only protospacers that are identical to the $WT$ plasmid.

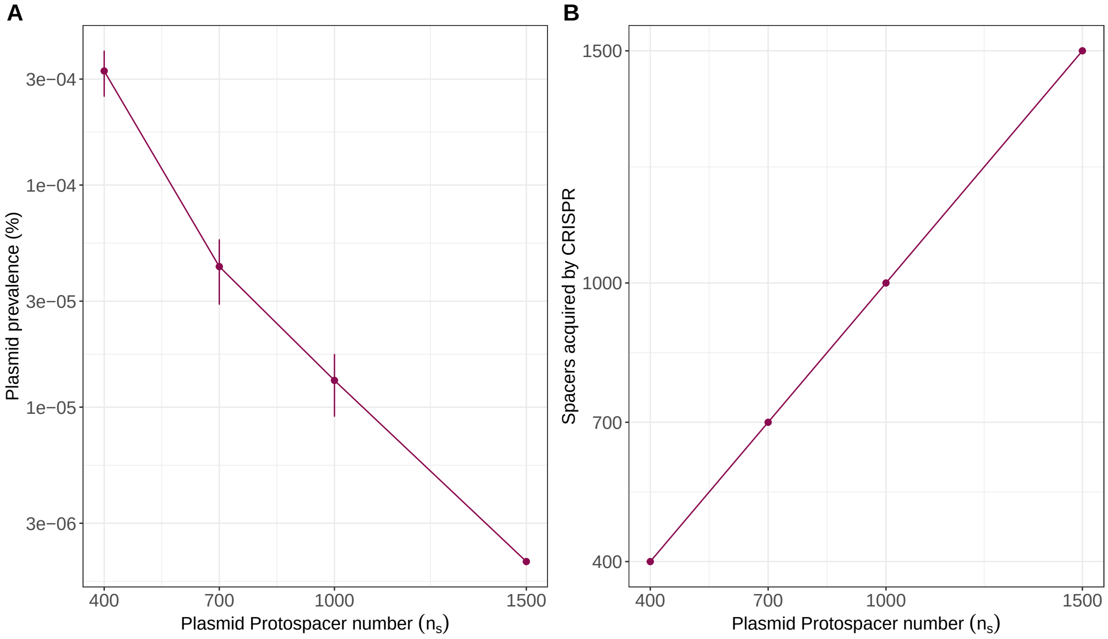


**Figure S11. Influence of the plasmid protospacer number $n_s$ on the plasmid prevalence and the number of different spacers acquired from CRISPR-Cas in the bacterial population. A.** Final plasmid prevalence decreases with higher plasmid protospacer number $n_s$. The points represent the mean prevalence for 5 simulations ran per value of $n_s$. The error bars represent the standard deviation. **B.** All available protospacers $n_s$ are acquired within the bacterial population by CRISPR. We set the values for the parameters as follows: $\alpha = 10^{-6}$, $\mu = 10^{-8}$, $\gamma_{wt} = \gamma_e = 10^{-10}$, $c_{wt} = 0.1$, $c_e = 0.2$, $\tau = 0.05$.

| | |
|---|---|
| $S$ | **Bacteria with naive CRISPR and no plasmid** |
| $I$ | **Bacteria with naive CRISPR and infected with wildtype plasmid** |
| $I_m$ | **Bacteria with naive CRISPR and infected with mutated plasmid** |
| $R$ | **Bacteria resistant against wildtype plasmid** |
| $R_{I_m}$ | **Bacteria resistant against wildtype plasmid and infected with mutated plasmid** |

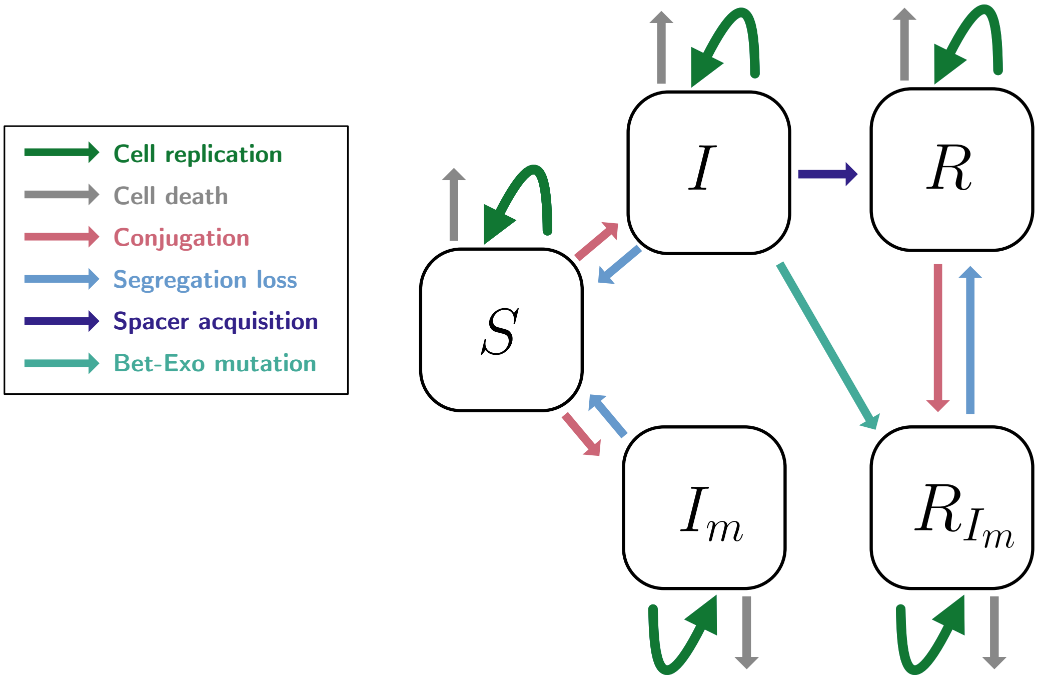


**Figure S12. Visualization of the modeled bacterial population, infection and immunity dynamics of CRISPR-Cas and a conjugative plasmid with *Bet/Exo*.**
